# Estimation of indirect genetic effects and heritability under assortative mating

**DOI:** 10.1101/2023.07.10.548458

**Authors:** Alexander Strudwick Young

## Abstract

Both direct genetic effects (effects of alleles in an individual on that individual) and indirect genetic effects — effects of alleles in an individual (e.g. parents) on another individual (e.g. offspring) — can contribute to phenotypic variation and genotype-phenotype associations. Here, we consider a phenotype affected by direct and parental indirect genetic effects under assortative mating at equilibrium. We generalize classical theory to derive a decomposition of the equilibrium phenotypic variance in terms of direct and indirect genetic effect components. We extend this theory to show that popular methods for estimating indirect genetic effects or ‘genetic nurture’ through analysis of parental and offspring polygenic predictors (called polygenic indices or scores — PGIs or PGSs) are substantially biased by assortative mating. We propose an improved method for estimating indirect genetic effects while accounting for assortative mating that can also correct heritability estimates for bias due to assortative mating. We validate our method in simulations and apply it to PGIs for height and educational attainment (EA), estimating that the equilibrium heritability of height is 0.699 (S.E. = 0.075) and finding no evidence for indirect genetic effects on height. We estimate a very high correlation between parents’ underlying genetic components for EA, 0.755 (S.E. = 0.035), which is inconsistent with twin based estimates of the heritability of EA, possibly due to confounding in the EA PGI and/or in twin studies. We implement our method in the software package *snipar*, enabling researchers to apply the method to data including observed and/or imputed parental genotypes. We provide a theoretical framework for understanding the results of PGI analyses and a practical methodology for estimating heritability and indirect genetic effects while accounting for assortative mating.

## Introduction

Since Galton’s 1886 work on the relationship between parent and offspring height^1^, explaining resemblance between relatives has been central to the biometrical approach to heredity. R.A. Fisher’s foundational 1918 paper, *The correlation between relatives on the supposition of Mendelian inheritance*^2^, unified the biometrical approach to heredity — whose intellectual lineage traces back to Galton — with Mendelian inheritance^3^. Fisher’s primary concern in this work was to show how Mendelian genetics, which describes the inheritance of discrete entities called alleles, could explain resemblance between relatives for continuous phenotypes like height. Fisher showed that, when alleles have additive effects, and there are no other sources of correlation between relatives, the phenotypic correlations between relatives are determined by the heritability of the phenotype, *h*^2^, and their coefficient of relatedness — provided that the population is infinite and mating randomly^2,4–6^.

After this groundbreaking result, Fisher’s paper goes on to address the more difficult problem of assortative mating (AM) — non-random mating leading to phenotypic correlation (usually assumed to be positive) between mothers and fathers, and therefore correlations between maternally and paternally inherited alleles. Fisher showed that AM induces a correlation between maternal and paternal genetic components, which increases the variance of the genetic component in the subsequent generation. The correlations between relatives’ genetic components and the variance of the genetic component increase towards equilibrium values as this process continues, requiring only a handful of generations to reach an approximate equilibrium^2,7–9^. Fisher derived the phenotypic correlations between relatives at equilibrium^2,4,6,7^, which Greg Clark showed can be closely fit to correlations between relatives’ social status in England in a dataset spanning 1600-2022 (ref^10^) — although these data do not rule out environmental effects of parents on offspring.

Building on Fisher’s work, Crow and Felsenstein gave an alternative derivation of the increase in genetic variance due to AM^11^, also detailed in Chapter 4 of Crow and Kimura’s textbook^7^. They showed that, for a phenotype affected by many genetic variants spread across the genome, the genetic variance is inflated by a factor of 1/(1 − *r*_*δ*_) at equilibrium, where *r*_*δ*_ is the correlation between maternal and paternal genetic components. Their method does not assume a particular model of AM, just that it has reached an equilibrium. The most common model of assortative mating states that all the correlation between parents’ genetic components is explained by matching on the observed phenotype, a model often called primary phenotypic assortment^12,13^. Assuming primary phenotypic assortment, and that the regression of phenotype onto genetic component is linear^8^, it can be shown that 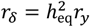, where 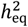 is the equilibrium heritability, and *r*_*y*_ is the correlation between parents’ phenotypes^2,7,8,11^.

The theory described above applies to phenotypes determined by additive effects of alleles in an individual on that individual, called direct genetic effects (DGEs), and random environmental effects/noise. In the 1970s, Cavalli-Sforza and Feldman developed models where, in addition to genetic transmission from parents to offspring, parental phenotypes affect the offspring’s phenotype through an environmental process called “vertical transmission” or “cultural transmission”^14,15^. The models of Cavalli-Sforza and Feldman were influential in the creation of the field of gene-culture coevolution^16^. In 1978, Cloninger, Rice, and Reich extended the models of Cavalli-Sforza and Feldman to include transmission of a general “cultural value” (possibly distinct from the offspring phenotype) from parent to offspring^17^. They extended this model to include AM due to matching on the phenotype, giving equilibrium results^18^. Their model makes predictions about the correlations between relatives^18^, and versions of their model have been used to analyse lifespan using a large pedigree from *Ancestry*.*com*^19^ and educational attainment (EA) using Swedish register data^20^. However, these analyses — like Clark’s analysis of social status in England^10^ — are unable to separate genetic transmission (heritability) from cultural transmission without making assumptions that are unlikely to be true.

Building on these vertical/cultural transmission models, behaviour genetics researchers extended the classical twin design — based on estimation of heritability by comparison of monozygotic (MZ) and dizygotic (DZ) twin pairs — to ‘extended twin and family designs’ (ETFDs) that also model the phenotypes of parents and other relatives of DZ and MZ twins. These ETFDs can model vertical transmission from parents, in addition to heritability^12^. The ‘stealth model’ from Truett et al. in 1994^21^ enabled modelling of assortative mating due to matching on the observed phenotype. Keller et al.^12^ introduced the ‘cascade model’ in 2009, a generalisation of the stealth model that allows the matching to take place on a latent, unobserved phenotype.

Although ETFDs can separately identify genetic and cultural transmission from parents while accounting for assortative mating, obtaining precise estimates of the parameters of these models can be difficult due to limited samples of twin pairs with phenotype data on their relatives^22^.

While empirical analyses of genotype-phenotype data have produced estimates of some important parameters that relate to vertical transmission models — including heritability^23–27^, correlations induced by assortative mating^13,28–30^, and indirect genetic effects (IGEs, also called ‘genetic nurture’)^31–33^ — robust estimation of vertical transmission model parameters remains challenging^22,33^.

IGEs are causal effects of alleles in one individual on another individual’s phenotype, mediated through the environment. When IGEs come from genetically related individuals, they contribute to the genotype-phenotype associations estimated in genome-wide association studies (GWASs) and lead to bias in heritability estimates from many methods^26,31,34,35^. This manuscript focuses on parental IGEs, effects of alleles in parents on their offspring through the rearing environment, but IGEs could come from other classes of relatives, such as siblings^31,34,36^, or from unrelated individuals. Vertical transmission models induce parental IGEs when the parental phenotype that affects offspring through the environment is heritable: the genetic variants that affect the parental phenotype will have IGEs on the offspring phenotype^33,34^. There is evidence that parental IGEs are important for educational outcomes^31,32^, but this evidence has been contested as confounded with the influence of population stratification and AM^34,37–39^.

AM generates bias in most methods for estimating heritability^23,40^, including: classical twin studies^23,41,42^; methods based on realized relatedness between relatives, such as siblings^23,24^ and more distant relatives, as in Relatedness Disequilibrium Regression, or RDR^23,26^; LD-score regression, or LDSC^40^; and genomic relatedness-matrix restricted maximum likelihood, or GREML^40^. While techniques for adjusting for the bias due to AM have been proposed^23^, they typically assume that all the correlation between maternal and paternal genetic components is explained by matching on the phenotype. This model has been shown to be inaccurate for EA, where the correlation between maternal and paternal genetic predictors of EA is far higher than can be explained by matching on the phenotype^13,20,43^. Factors that may contribute to maternal and paternal genetic predictors (or components) becoming more correlated than expected due to matching on the observed phenotype include: matching on a correlated phenotype that is more highly correlated with the underlying genetic predictor/component than the observed phenotype^12,13,30^, and matching based on the phenotypes of the mate’s family members — and/or ancestry — in addition to the phenotype of the mate. It would therefore be desirable to have a technique for adjusting for bias due to AM that does not assume the correlation between parents’ genetic components is entirely due to matching on the phenotype.

Most studies examining evidence for IGEs have proceeded by correlating parental alleles not transmitted to offspring with offspring phenotypes^31,32^. While this correlation captures IGEs, it also partly captures the genetic component of the phenotype with which the non-transmitted alleles are correlated due to AM^31,33,38,39^. Kong et al.^31^ attempted to adjust for this bias due to AM, concluding the bias was small. However, they did not measure the uncertainty in their adjustment, and the adjustment relied upon assuming that there had been only one generation of AM and that DGEs and IGEs are perfectly correlated. Balbona et al. proposed a structural equation model (SEM) to adjust for the bias due to AM in estimates of IGEs^33^, but this method assumes correlations between parents’ genetic components are due entirely to matching on the phenotype. The method of Balbona et al. additionally requires observations on the parental phenotype through which IGEs/vertical transmission operate, or knowledge of the true heritability of the phenotype and an assumption that the parental phenotype through which IGEs/vertical transmission operates is the same as the offspring phenotype.

In this paper, we generalize Crow and Felsenstein’s^11^ approach (which considered only DGEs) to also include parental IGEs. We derive a decomposition of the equilibrium phenotypic variance in terms of DGE and IGE components that does not assume a particular model of assortative mating. We connect our results to estimation of heritability using random-variation in realized relatedness due to Mendelian segregations^24,26^, deriving the bias in these methods due to AM. We extend our approach to derive results on analysis of polygenic predictors (called polygenic indices or PGIs — also known as polygenic scores). We show how to assess evidence for IGEs from PGI analysis when there is AM and to adjust for bias in heritability estimates due to AM. We apply these results to PGIs for height and EA, and we correct for the bias in the height heritability estimate due to AM.

## Results

### Phenotype model

We model the phenotype of sibling *j* in family *i* as the result of DGEs, IGEs from parents, and a residual environment/noise term:

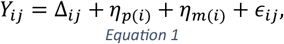

where

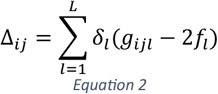

is the DGE component; *δ*_*l*_ is the direct effect of variant *l*; *g*_*ijl*_ is the genotype of sibling *j* in family *i* at variant *l*; and variants are assumed to be bi-allelic with frequency *f*_*l*_, constant across generations, so that *E*[*g*_*ijl*_] = 2*f*_*l*_. The paternal and maternal IGE components are

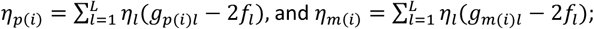

where *η*_*l*_ is the (average) parental indirect genetic effect of variant *l*; and *g*_*p*(*i*)*l*_ and *g*_*p*(*i*)*l*_ are, respectively, the genotypes of father and mother in family *i* at variant *l*. We model only the average parental IGE here, as, for the results in this manuscript, differences between paternal and maternal IGEs can be subsumed into the residual and ignored (Supplementary Note Section 3.4).

AM induces a correlation between parents’ DGE components, which we define to be:

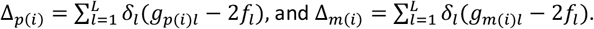

We give a glossary of terms used in the paper after the Discussion.

### Equilibrium phenotypic variance

We decompose the equilibrium phenotypic variance in terms of the random mating variance decomposition and the equilibrium correlations between parents’ DGE and IGE components. Figure 1 shows the notation for the equilibrium correlations: the within-parent (or cis-parental) correlation, 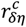, between DGE and IGE components are the same for mothers and fathers because the DGEs (*δ*_*l*_) and (average) IGEs (*η*_*l*_) do not depend on the parent; similarly for the cross-parent (or trans-parental) correlation, 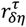. To make the decomposition tractable, we assume that the *L* causal variants segregate independently, implying they would be uncorrelated in a random mating population.

**Figure 1.**
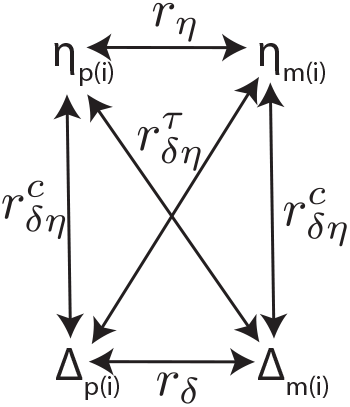
Diagram of correlations between parents’ direct genetic effect (DGE) and indirect genetic effect (IGE) components. Δ_p(i)_ and Δ_m(i)_ are the paternal and maternal DGE components with correlation r_δ_. η_p(i)_ and η_m(i)_ are the paternal and maternal IGE components with correlation r_η_. The within-parent (or cis-parental) correlation between DGE and IGE components is 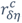, for which we use a superscript ‘c’ to denote ‘cis-parental’. The cross-parent (or trans-parental) correlation between DGE and IGE components is 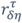, for which we use a superscript ‘τ’ to denote ‘trans-parental’.

The random-mating variance decomposition is the same as given in Young et al. 2018, who derived it for the RDR method for estimating heritability^31^:

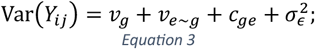

where 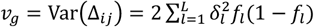 is the random-mating genetic variance; 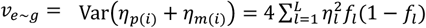 is the random-mating variance due to (average) parental IGEs; and 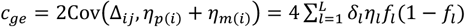 is the random-mating variance due to covariance between DGEs and parental IGEs; and 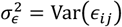.

In Supplementary Note Section 3, we show that, for AM at equilibrium and in the limit as *L* → ∞,

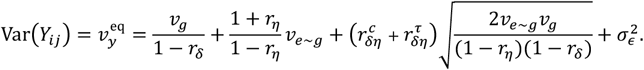

We define the equilibrium variance components:

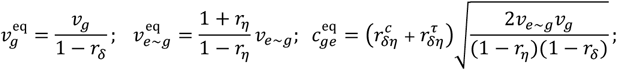

and the equilibrium heritability, 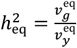.

The equilibrium phenotypic variance due to covariance between DGE and IGE components, 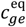, will be non-zero when 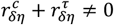. This is true even when DGEs and IGEs are uncorrelated but AM induces a correlation between DGE and IGE components 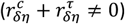. In other words, AM induces a correlation between the DGE and IGE components even if there would be none under random mating.

In theory, it would be possible to estimate this variance decomposition given estimates of the random-mating variance components and correlations between parents’ DGE and IGE components (Figure 1). In Supplementary Note Section 3.4, we show that differences between maternal and paternal IGEs do not alter the equilibrium variance decomposition: the variance component due to parental IGE asymmetry is absorbed into the residual and is unchanged by AM. Furthermore, since proband PGI and average parental PGIs are uncorrelated with the component due to parental IGE asymmetry^34^, such asymmetries do not affect the PGI analysis results contained in this manuscript.

If *c*_*ge*_ ≠ 0 (DGE and IGE components are correlated under random mating) then the equilibrium variance decomposition can be expressed in terms of *c*_*ge*_ (Supplementary Note Section 3.3.1):

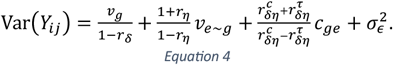

In Supplementary Note Section 3.3.2, we show that

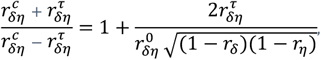

where 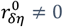 is the random-mating correlation between DGE and IGE components, which equals the genome-wide correlation between standardized DGEs and IGEs. This shows that AM will increase the magnitude of the variance due to covariance between DGE and IGE components provided that 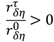.

If we assume primary phenotypic assortment, then it can be shown that (Supplementary Note Section 3.5):

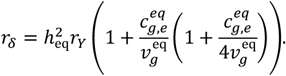

Thus, the correlation between parents’ DGE components can differ from 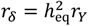 (the classic result for primary phenotypic assortment^7^) when there is (potentially AM induced) correlation between DGE and IGE components. When DGE and IGE components are positively correlated, the equilibrium correlation between parents’ genetic components will thus be higher than would be predicted using the classic result, 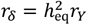, and consequently the AM induced inflation of phenotypic variance due to DGEs would be larger.

### Estimating heritability using realized relatedness

While siblings have a relatedness coefficient of ½ in expectation — based on the expected proportions of the genome shared identical-by-descent (IBD) from each parent — there is variation around this expectation due to random segregation of genetic material in the parents during meiosis. The realized relatedness between siblings is computed from the proportions of the genome shared IBD from each parent, and therefore captures the random variation in relatedness around the expectation. In outbred samples, the realized relatedness between siblings has an approximate normal distribution with mean close to 0.5 and a standard deviation around 0.04 (ref^24,26^). By examining how the phenotypic correlation between siblings changes with realized relatedness, an estimate of heritability can be obtained that is robust to population stratification^24,26^. We call this method ‘sib-regression’.

Here we examine how realized relatedness affects the phenotypic correlation between siblings in our model with DGEs, IGEs, and AM at equilibrium. In Supplementary Note Section 4, we show that, for a sibling pair with phenotypes (*Y*_*ij*_, *Y*_*ik*_) and realized relatedness *R*_*ijk*_,

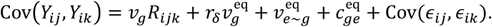

(This is the result for the limit as the effective number of independent loci contributing to the DGE component goes to infinity. We give results for a finite number of loci in Supplementary Note Section 4.) This shows that variation in realized relatedness gives information about the random mating variance of the DGE component, not the equilibrium variance, because the variation in realized relatedness is due to random segregation of genetic material in a family where the correlations induced by AM are irrelevant.

Heritability can be estimated by a regression of 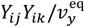 onto *R*_*ijk*_ across sibling pairs — where we have assumed the phenotypes have mean zero. The slope of this regression gives the estimate of heritability. Assuming that the realized relatedness is uncorrelated with *∈*_*ij*_*∈*_*ik*_ — which would be violated when there are IGEs between siblings^26^ — we show in Supplementary Note Section 4 that the slope of the regression is 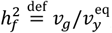, the random-mating variance of the DGE component variance divided by the equilibrium phenotypic variance. The estimand 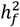 is smaller than the equilibrium heritability, 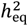, by a factor of (1 − *r*_*δ*_). Thus, one could estimate 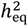 by inflating estimates of 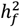 by a factor of 1/(1 − *r*_*δ*_).

The intercept of the regression is 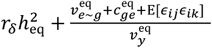. This includes a term, 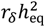, that will be non-zero when the phenotype is heritable and there is AM, even in the absence of IGEs or other environmental effects shared between siblings. This implies that, when there is AM, the intercept will give an upward biased estimate of the proportion of phenotypic variance explained by environmental effects shared between siblings^24^. To correct for the bias due to AM, one could subtract an estimate of 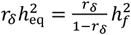 from the intercept of the regression.

Our theoretical results for AM at equilibrium agree with Kemper at al., who argued that sib-regression estimates the random mating genetic variance divided by the phenotypic variance in the present generation^23^, which is the equilibrium phenotypic variance at equilibrium, as in our model/derivation. Kemper et al. supported their argument with simulations of a single generation of AM and a theoretical derivation that, although it reached the correct conclusion, is invalid (Supplementary Note Section 4.2). Kemper et al. argued that RDR^26^, which is a generalization of sib-regression to all relative pair classes, also estimates 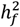. Our theoretical results imply that this is true since RDR — like sib-regression — uses within-family variation in realized relatedness to estimate heritability.

Although they do not use realized relatedness, classical twin studies based on MZ-DZ twin comparisons have also been shown to estimate 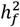 under the ACE model when AM is at equilbrium^23^. It is trivial to show the same result holds in our model. This implies that heritability estimates from the ACE twin model, from sib-regression, or from RDR can be combined with estimates of *r*_*δ*_ to estimate the equilibrium heritability.

In the following sections, we show how analysis of genetic predictors (called polygenic indices, or PGIs; also called polygenic scores, or PGS) can be used to estimate *r*_*δ*_, to adjust heritability estimates for bias due to AM, and to assess evidence for parental IGEs while accounting for AM.

### Family-based polygenic index analysis

Building on Kong et al.^31^, many studies have examined both offspring and parental PGIs as predictors of offspring phenotypes^32,34,43^. In the following sections, we show how to interpret the results of these studies in our model.

A PGI is a weighted sum of genotypes (numbers of copies of alleles) across genetic variants:

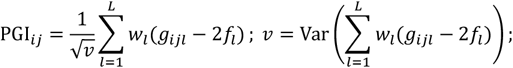

where PGI_*ij*_ is the PGI of sibling *j* in family *i*, and 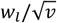 is the weight of variant *l*. If we set *w*_*l*_ = *δ*_*l*_ for all loci, then PGI_*ij*_ ∝ Δ_*ij*_, as defined in Equation 2. In this section, we assume the PGI has been normalized to have variance 1, i.e. the un-standardized PGI has been divided by its standard deviation, 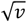, where *v* is its variance. Under random-mating,

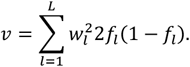

We now define the paternal and maternal PGIs using the same weights:

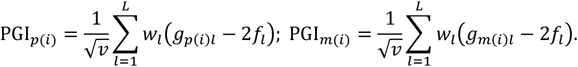

As they use the same weights, the maternal and paternal PGIs have the same variance as the offspring PGIs under random mating and under AM at equilibrium.

Given parental genotypes, offspring genotypes vary due to random segregations during meiosis in the mother and father. Furthermore, Mendelian inheritance induces an important relationship between parent and offspring PGIs. Letng *G*_par(*i*)_ represent the genotypes of the parents:

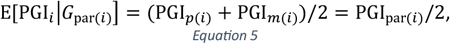

where PGI_par(*i*)_ = PGI_*p*(*i*)_ + PGI_*m*(*i*)_. This result holds generally, whether there is AM or not.

The most common type of PGI analysis is a regression of phenotype onto PGI without controlling for parental PGIs (but potentially controlling for covariates, such as genetic principal components):

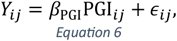

where *β* is called the ‘population effect’ of the PGI as it reflects the overall association of the PGI and phenotype in the population (aNer accounting for covariates).

Most of the evidence for parental IGEs has derived from fitting a version of the following regression equation^31,32,43^:

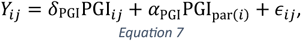

where *δ*_PGI_ is called the direct genetic effect of the PGI, and *α*_PGI_ is called the average non-transmitted coefficient (NTC), as it reflects the correlation of offspring phenotype with the PGI constructed from the parental alleles not transmitted to the offspring^34^. Often, *α*_PGI_ has been interpreted as reflecting IGEs alone, which would be true under random-mating, but it can also reflect population stratification and, as we detail below, AM^34,38,39^. Since offspring PGI is conditionally independent of environment given parental genotypes, and E[PGI_*i*_|*G*_par(*i*)_] = PGI_par(*i*)_/2 (Equation 5), *δ*_PGI_ reflects DGEs of causal variants alone, and does not include IGEs or other forms of gene-environment correlation, e.g. population stratification^31,34^.

In Supplementary Note Section 5, we show that, assuming random-mating,

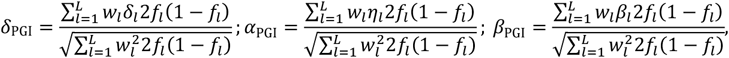

where *β*_*l*_ = *δ*_*l*_ + *η*_*l*_. In Table 1, for certain special values of the weight vector *w*_*l*_, we give *δ*_PGI_, *α*_PGI_, and *β*_PGI_ in terms of the random-mating variance components^26^ (Equation 3), and the random-mating correlation between DGE and IGE components, 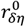.

**Table 1.** Expected regression coefficients for two-generation PGI analysis under random-mating. Direct effect (δ_PGI_), average non-transmitted coefficient (α_PGI_), and population effect (β_PGI_) for standardized PGIs with different weight vectors, specified by the w_l_ column, which gives the weight for variant l in the un-standardized PGI. Here, we give the regression coefficients for the PGI standardized to have variance 1. The PGI coefficients are expressed in terms of the random-mating variance components (Equation 1). The variance explained by the population effect PGI under random mating is 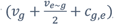, which is also the variance estimated by GREML applied to all causal SNPs in a random mating population^26^. These results are for idealized weight vectors given by the true DGEs, δ_l_, average parental IGEs, η_l_, and population effects, β_l_ = δ_l_ + η_l_. These results are not valid when weights are estimated with noise or bias — as would be the case for real world PGIs computed from GWAS summary statistics — or when there is non-random mating. A glossary of symbols is included after the Discussion.

| PGI | $w_l$ | $\delta_{\text{PGI}}$ | $\alpha_{\text{PGI}}$ | $\beta_{\text{PGI}}$ |
| --- | --- | --- | --- | --- |
| DGE | $\delta_l$ | $\sqrt{v_g}$ | $r_{\delta\eta}^0 \sqrt{\frac{v_{e\sim g}}{2}}$ | $\sqrt{v_g} + r_{\delta\eta}^0 \sqrt{\frac{v_{e\sim g}}{2}}$ |
| Average Parental IGE | $\eta_l$ | $r_{\delta\eta}^0 \sqrt{v_g}$ | $\sqrt{\frac{v_{e\sim g}}{2}}$ | $\sqrt{\frac{v_{e\sim g}}{2}} + r_{\delta\eta}^0 \sqrt{v_g}$ |
| Population Effect | $\beta_l$ | $\frac{v_g + c_{g,e}/2}{\sqrt{v_g + v_{e\sim g}/2 + c_{g,e}}}$ | $\frac{(v_{e\sim g} + c_{g,e})/2}{\sqrt{v_g + v_{e\sim g}/2 + c_{g,e}}}$ | $\sqrt{v_g + v_{e\sim g}/2 + c_{g,e}}$ |

### Two-generation analysis of the direct genetic effect PGI at equilibrium

We now give results for PGI analysis under AM at equilibrium. First, we give results for analyzing the true DGE PGI, as results for DGE PGIs that do not capture all the heritability can be expressed in terms of coefficients for the true DGE PGI. We define (Equation 2):

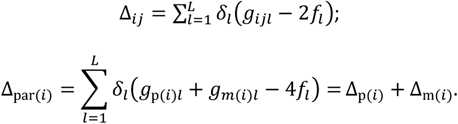

Consider analyzing this PGI in a two-generation regression model:

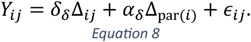

In Supplementary Note Section 6, we show that, at equilibrium:

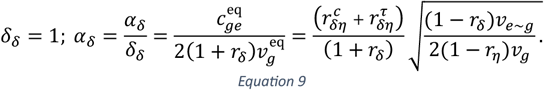

This shows that we expect the average NTC of the DGE PGI to be non-zero only when IGEs are present and 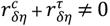, which will almost surely be true when there is AM, even when DGEs and IGEs are uncorrelated. To obtain equivalent results for the true DGE PGI standardized to have variance 1, we multiply both *δ*_*δ*_ and *α*_*δ*_ by 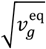. Let 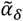 be the average NTC for the standardized true DGE PGI, then

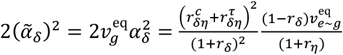

This implies that 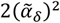 gives a measure proportional to the variance explained by parental IGEs, provided that 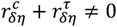. In the random-mating case, this will be less than *v*_*e*∼*g*_ unless 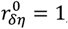, i.e. unless the genome-wide correlation between DGEs and IGEs is 1. While this captures part of the variance explained by parental IGEs, it does not capture the phenotypic variance due to covariance between DGE and IGE components. In Supplementary Note Section 6.1, we show that the proportion of phenotypic variance explained by joint regression onto the parental and offspring DGE PGIs (Equation 8) is:

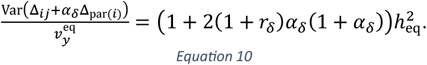

If there are no parental IGEs, or the IGE component is uncorrelated with the DGE component, then *α*_*δ*_ = 0, and the proportion of variance explained by parental and offspring DGE PGIs would be 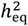. Thus, the contribution of parental IGEs to the variance explained by the parental and offspring DGE PGIs (due to both the IGE component and its covariance with the DGE component) is

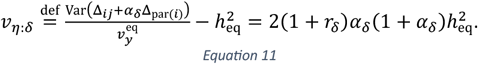

Thus, the contribution from parental IGEs is magnified when *r*_*δ*_ > 0, i.e. when (positive) AM is present.

This shows that the true DGE PGI — which could be estimated from family-based GWAS^34,44^ — can be used to assess the contribution of parental IGEs to the phenotypic variance. However, when the estimated DGE PGI does not capture all the heritability and there is assortative mating, the situation is more complicated, and requires a different solution, as we outline below.

### Two-generation analysis of an incomplete direct genetic effect PGI at equilibrium

In real-world applications, we will never have access to the true DGE PGI. Even with unbiased estimates of DGEs, sampling errors due to finite sample size^45^ mean that any real-world DGE PGI will fail to capture all of the heritability. Differences in local linkage disequilibrium patterns and/or DGEs between training data and target data^39,46,47^ would further reduce the heritability explained by a real-world PGI. If using population effect estimates from standard GWAS — as is currently standard practice — bias due to IGEs and improperly controlled population stratification^34^ mean that such PGIs are unlikely to capture all of the heritability even as sampling error approaches zero. Furthermore, incomplete genotyping and/or imperfect imputation of rare variants and structural variants^48–50^ mean that not all relevant genetic variation is captured by current PGIs.

To accommodate the complexities of real-world applications as best we can, we develop theory for a DGE PGI that explains a fraction, *k*, of the heritability in a random-mating population. The model we use assumes that we include only a fraction, *k*, of the independently segregating causal variants, which are assumed to have equal frequency and equal effect size, *δ*:

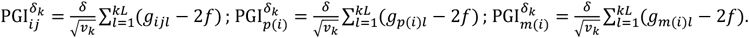

We define the PGIs such that they have been standardized to have variance 1 through division by 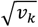, where 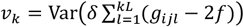, which is assumed to be the same for parent and offspring PGIs because we are assuming equilibrium.

Although this simplified model is not realistic and does not capture all the possible ways a PGI may capture only a fraction, *k*, of the heritability in a random-mating population, we show that the results hold under more general conditions through simulations (below).

For the following sections, we assume we have a sample of families where we have observed the phenotypes of the offspring, *Y*_*ij*_, along with the offspring and parental incomplete DGE PGIs: 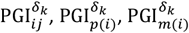. We denote the correlation between maternal and paternal incomplete DGE PGIs to be

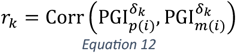

We consider what we can learn from performing a two-generation PGI analysis using the incomplete DGE PGI. Specifically, consider we have performed a regression of the standardized offspring phenotype onto standardized offspring and parental PGIs:

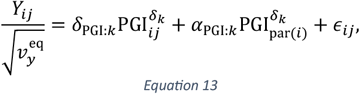

and let 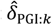 and 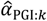 be the resulting estimates of the direct effect, *δ*_PGI:*k*_, and average NTC, *α*_PGI:*k*_, of the incomplete DGE PGI.

We also consider estimating the population effect of the incomplete DGE PGI, *β*_PGI:*k*_, by the following regression:

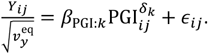

Because 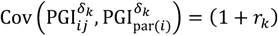 at equilibrium, it is trivial to show that

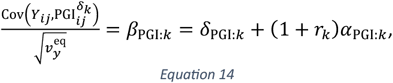

which gives a useful connection between the results of one and two-generation PGI analyses.

### Impact of assortative mating on PGI analysis in the absence of indirect genetic effects

AM can induce statistical properties that can be confused with the influence of IGEs. In Supplementary Note Section 7.3.1, we show that the fraction of phenotypic variance the incomplete PGI explains at equilibrium in a model without IGEs is:

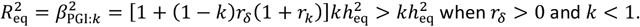

In other words, AM inflates the *R*^2^ between phenotype and PGI by a factor of 1 + (1 − *k*)*r*_*δ*_ (1 + *r*_*k*_). As *k* → 1, the inflation tends to zero. This is because the inflation is due to the correlation between the PGI and the DGE component that the PGI would be uncorrelated with in a random-mating population but becomes correlated with due to AM: when *k* = 1, there is no residual DGE component to be correlated with.

The AM-induced correlation between the PGI and the DGE component the PGI would be uncorrelated with in a random mating population affects results from two-generation PGI analysis. Since Kong et al.^31^, attention has been given to the ratio between direct and population effects (*δ*_PGI_/*β*_PGI_) for PGIs, since this measures how much apparent PGI effects ‘shrink’ when estimated within-family^43,51^ — a statistical signature of IGEs. However, population stratification and AM can also lead to shrinkage of PGI effects within-family^33,39,43,52^, implying that IGEs cannot be identified from ‘shrinkage’ of PGI effects within-family alone.

In Supplementary Note Section 7.3, we show that, for a PGI that would explain a fraction *k* of the heritability in a random mating population, this ratio is

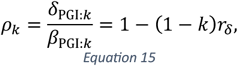

at equilibrium in the absence of IGEs (and no population stratification, which we do not include in our model). This equation thus gives a baseline expectation for what ‘shrinkage’ to expect based purely on AM. The equation shows that we could expect substantial shrinkage when *k* is not close to 1, and *r*_*δ*_ is substantially above 0 (Figure 2).

**Figure 2.**
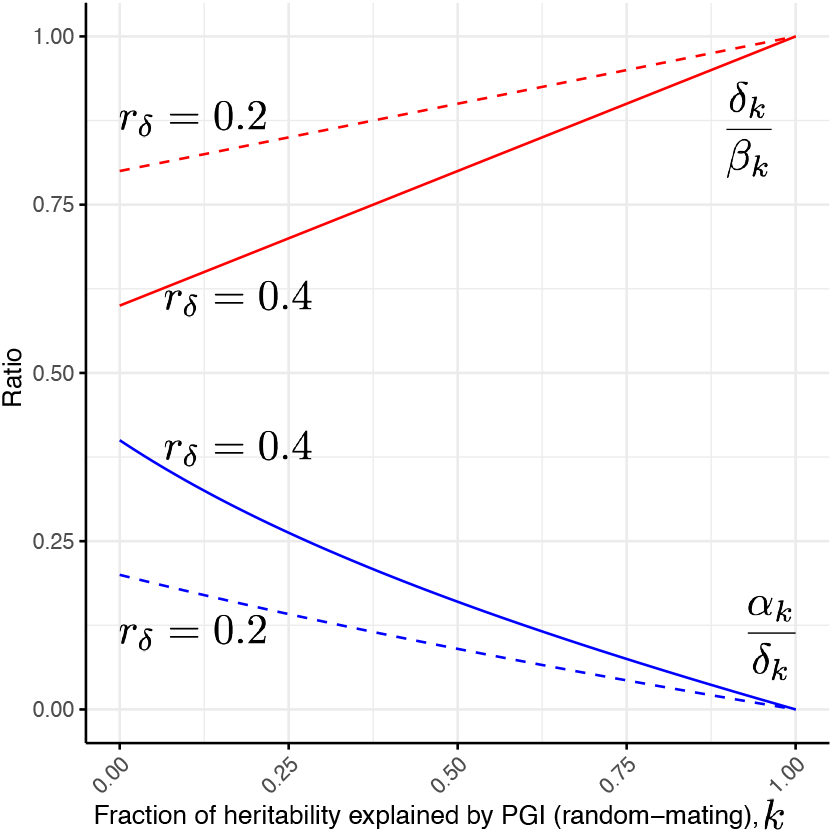
Impact of assortative mating on PGI analysis. We give results for the expected regression coefficients in one- and two-generation PGI analysis (Equations 13 and 14) under assortative mating at equilibrium without indirect genetic effects. The y-axis gives ratios: between the average NTC and direct effect, α_k_/δ_k_, in blue; and between direct and population effects, ρ_k_ = δ_k_/β_k_, in red. We plot these ratios as a function of the fraction of heritability the PGI would explain in a random-mating population, k, on the x-axis. As these ratios (Equations 15 and 17) also depend upon the correlation between parents’ direct effect components, r_δ_, we show the ratios as a function of k (x-axis) for both r_δ_ = 0.4 (solid lines) and r_δ_ = 0.2 (dashed lines). A glossary of symbols is included after the Discussion.

Consider analysing a single variant that explains a negligible amount of the heritability, then *ρ*_*k*_ = lim_*k*→0_(1 − (1 − *k*)*r*_*δ*_) = 1 − *r*_*δ*_. Versions of this result for a single variant has been given before by several authors^8,28,52^. The ratio 1 − *r*_*δ*_ gives the expected ‘shrinkage’ when estimating the DGE of a variant in family-based GWAS compared to estimating the population effect using standard GWAS. This raises the possibility that *r*_*δ*_ could be estimated from the average ‘shrinkage’ of DGEs compared to population effects of genome-wide SNPs, assuming AM is the only source of shrinkage.

### Estimating indirect genetic effects accounting for assortative mating

The above section suggests that one way to assess evidence for IGEs while accounting for AM would be to compute the ratio between direct and population effects for a particular PGI, and to compare this to an estimate of *ρ*_*k*_ (Equation 15), 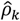, the ratio that would be expected due to AM alone without IGEs. If the estimated ratio between direct and population effects, 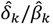, was statistically significantly different from 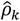, then this would constitute evidence that IGEs (and/or other forms of gene-environment correlation, such as population stratification) are present.

In the Methods and Supplementary Note Sections 7-9, we derive a similar but more formal procedure for performing two-generation PGI analysis accounting for AM (Figure 3). The inputs of this procedure are: an estimate of the correlation between parents’ incomplete DGE PGIs, 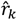; estimated regression coefficients from a regression of standardized offspring phenotype onto standardized offspring and parental (incomplete) DGE PGIs (Equation 13), 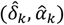; and an estimate of 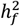, such as from MZ-DZ twin comparisons (ACE model), RDR, or sib-regression. These inputs are first used to estimate 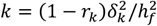, the fraction of heritability the PGI would explain in a random-mating population (Methods and Supplementary Note Section 8). Estimates of *k* and *r*_*i*_ can then be combined to estimate *r*_*δ*_ using a relationship between *r*_*δ*_ and *r*_*k*_ we derive (Supplementary Note Section 7.2):

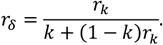

**Figure 3.**
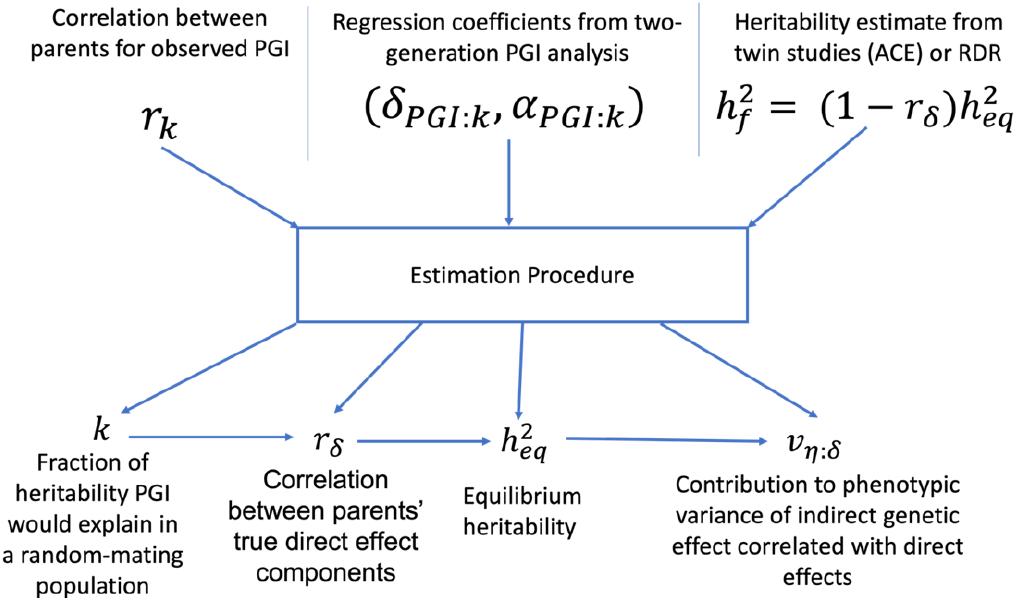
Schematic of two-generation PGI analysis accounting for assortative mating. A PGI is used as an instrument in order to make inferences about the impact of assortative mating (AM) and indirect genetic effects (IGEs) on phenotype variation. The inputs are the correlation between parents’ observed PGIs, r_k_; the regression coefficients from two-generation PGI analysis (Equation 13); and an estimate of 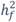 from MZ-DZ twin comparisons (ACE model), sib-regression, or RDR. These inputs are then put through a series of non-linear estimating equations (Methods) in order to estimate k, the fraction of heritability the PGI would explain in a random-mating population; r_δ_, the correlation between parents’ true direct genetic effect (DGE) components; 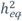, the equilibrium heritability (which is larger than 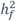 when there is AM); and v_η: δ_ the proportion of phenotypic variance contributed by the IGE component that is correlated with the DGE component (when a PGI constructed from unbiased DGE estimates is used). A glossary of symbols is included after the Discussion.

Given *r*_*δ*_ and 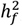, the equilibrium heritability is given by 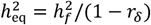. The IGE of the true DGE PGI, *α*_*δ*_ (Equation 9), can be estimated two ways. One is based on the ratio between direct and population effects of the incomplete DGE PGI, which we show to be (Supplementary Note Section 7.4):

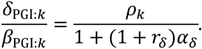

By rearranging, we obtain an expression for *α*_*δ*_:

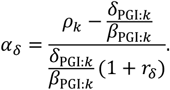

This equation matches the intuition that if the direct to population effect ratio, 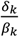,is smaller than would be predicted under a model without IGES (*ρ*_*k*_), this implies *α*_*δ*_ > 0, i.e. that there is an IGE component positively correlated with the DGE component.

We then estimate *α*_*δ*_ using estimates of *δ*_*k*_, *β*_*k*_, *k, r*_*δ*_, and *ρ*_*k*_ (Methods):

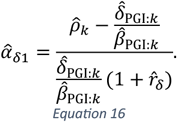

An alternative route to estimating *α*_*δ*_ is to use the ratio between the average NTC and the direct effect of the PGI: *α*_PGI:*k*_/*δ*_PGI:*k*_. Kong et al. estimated that this ratio was 0.427 for an EA PGI, and used this as the basis of their argument that IGEs on EA are substantial^31^. In Supplementary Note Section 7.4, we show that:

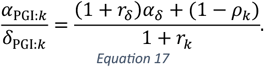

This shows that the average NTC is the sum of two components: one due to AM-induced correlation with the DGE component that the PGI would be uncorrelated with under random mating, (1 − *ρ*_*k*_)/(1 + *r*_*k*_); and one due to parental IGEs, (1 + *r*_*δ*_)*α*_*δ*_/(1 + *r*_*k*_). By rearrangement and substitution, we obtain:

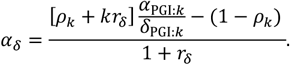

This yields a second sample estimator of *α*_*δ*_:

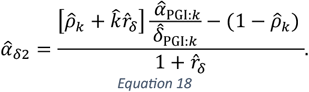

Although less intuitive than the estimator based on the ratio between direct and population effects (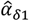, Equation 16), 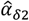 is generally to be preferred because 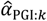 and 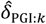 can be estimated from the same regression (Equation 13), making it easier to compute the approximate sampling variance (Supplementary Note Section 9). Simulations indicated that, when estimated from the same data, 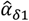 and 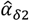 give almost identical results (Supplementary Table 7).

Given an estimate of *α*_*δ*_, 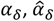, along with an estimate of 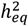 (above), one can then estimate the fraction of phenotypic variance contributed by the IGE component that is correlated with DGE component, *v*_*η*: *δ*_ (Equation 11). We estimate this as:

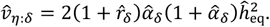

We implemented the estimation procedure (Figure 3) in *snipar* (*https://github.com/AlexTISYoung/snipar*). By inputting an estimate of 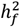 along with the data required to perform two-generation PGI analysis (offspring phenotypes and genotypes/PGIs, and observed and/or imputed parental genotypes/PGIs), *snipar* will estimate *δ*_PGI:*k*_, *α*_PGI:*k*_, *β*_PGI:*k*_, *r*_*k*_, *k, r*_*δ*_, 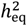, *ρ*_*k*_, *α*_*δ*_, and *v*_*η*: *δ*_, along with approximations to their sampling variances (Methods and Supplementary Note Section 9).

### Simulation study

We simulated 16 phenotypes with varying parameters using the *simulate.py* module in *snipar* (Methods and Supplementary Table 1). For each phenotype, we simulated the first-generation by random mating. We simulated 30,000 independent families and 1,000 causal SNPs, with two full-sibling offspring in each family. We simulated a DGE component that explained 50% of the phenotypic variance, i.e. *v*_*g*_/*v*_*y*_ = 0.5, in the first-generation. For some phenotypes, we also simulated a parental IGE component that explained 12.5% of the variance in the first generation, i.e. *v*_*e*∼*g*_/*v*_*y*_ = 0.125. We simulated DGEs and IGEs of individual SNPs from a bivariate normal distribution. We set the correlation between DGEs and IGEs, 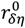, to 0, 0.5, or 1. For each set of IGE parameters, we simulated phenotypes affected by AM of varying strengths: the phenotypic correlation between parents in each generation was set to 0, 0.25, 0.5, and 0.75. In order to reach approximate equilibrium, we simulated 20 generations of mating after the first-generation produced by random mating.

We found a close agreement between our theoretical results on the equilibrium phenotypic variance decomposition (Equation 4) and the simulation results (Supplementary Table 1 and Supplementary Figure 1). Using the last two simulated generations, we performed two-generation PGI analysis using PGIs constructed from the true DGEs plus estimation error (Methods and Supplementary Tables 2-5). Although the theory was derived assuming that we have a PGI constructed using the true DGEs as weights for a subset of causal variants, we tested a different scenario, in which we used the true DGEs plus estimation error (as could be obtained from a family-based GWAS) as weights:

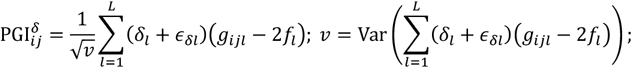

where *∈*_*δl*_ were simulated as independent variables with *N*(0, *v*_*∈δ*_) distribution. The variance of the estimation error, *v*_*∈δ*_, was set in multiples of the variance of the true DGEs: 0 (standardized true DGE PGI), 1, 10, and 100. The fraction of heritability explained by the PGIs in a random-mating population (*k*) was thus approximately equal to 1/(1 + *v*_*∈ δ*_). So, for *v* = 0, 1,10,100, *k* ≈ 1, 0.5, 0.09, 0.01.

To complete the inputs (Figure 3), we estimated *r*_*k*_ using the sample correlation between the parents’ PGI values, and we used the true value of 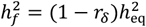 as the heritability input. We give results for *v* = 1, *k* ≈ 0.5 in Figure 4 and Supplementary Table 3, which shows the procedure produces approximately unbiased estimates of *r*_*δ*_, 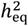, *α*_*δ*_, and *v*_*η*: *δ*_ with accurate standard errors. However, while the inference procedure produces accurate results when *k* ≈ 0.5, and reasonably accurate results for *k* ≈ 0.09 (Supplementary Figure 2 and Supplementary Table 4), it produces biased and unstable results when *k* ≈ 0.01 (Supplementary Table 5 and Figure 5b). This is because, when *δ*_*k*_, *α*_*k*_, and *r*_*k*_ are close to zero, noise in the estimates makes the inference unstable since many terms in the estimating equations involve ratios of parameters.

**Figure 4.**
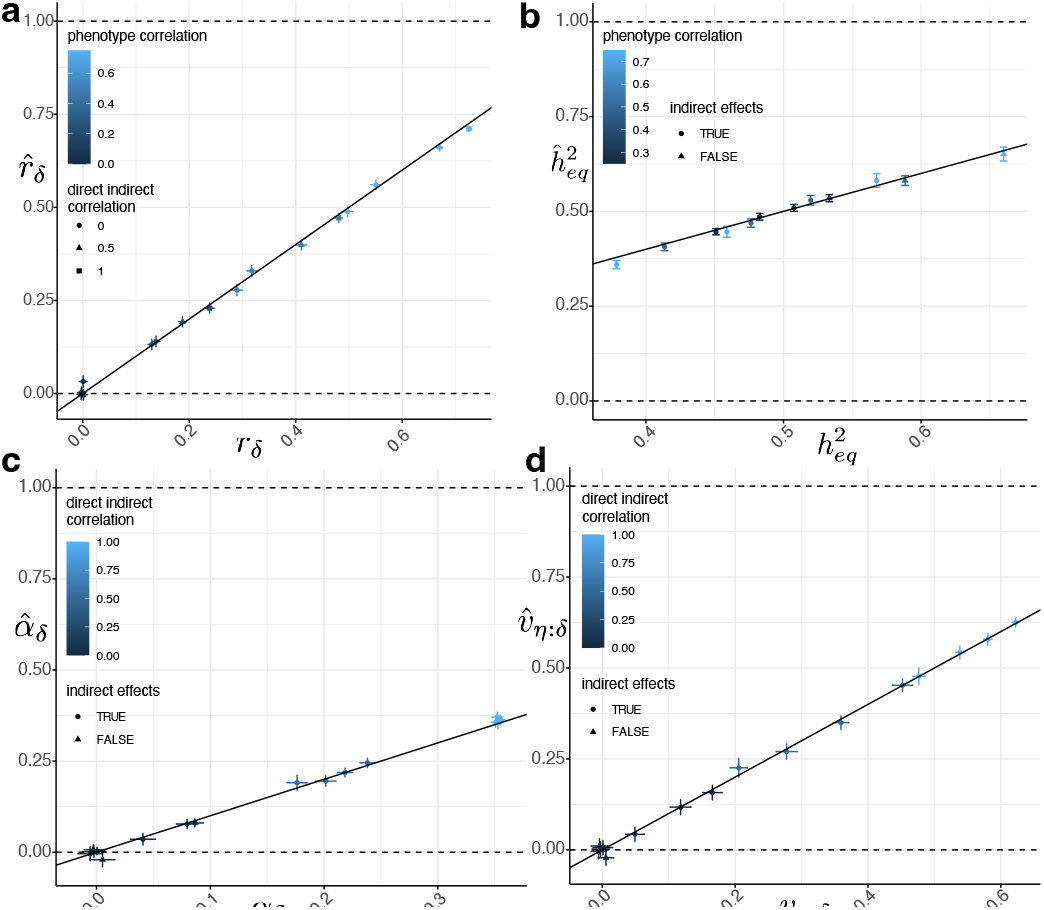
Simulation results for a direct genetic effect PGI. Across 16 simulated phenotypes (Methods), we computed PGIs using weights equal to the true direct genetic effects (DGEs) plus a noise term of variance equal to the variance of the true DGEs, simulating estimation error. This gave a DGE PGI that explained approximately 50% of the heritability in a random-mating population (Supplementary Table 3). We performed two-generation PGI analysis (Methods and Figure 3) in order to estimate a) r_δ_, the correlation between parents’ true DGE components (that explain all of the heritability); b) 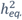, the equilibrium heritability; c) α_δ_, the indirect genetic effect (IGE) of the true DGE PGI; and d) the proportion of phenotypic variance contributed by the IGE component that is correlated with the DGE component, v_η: δ_. Vertical and horizontal error bars indicate 95% confidence intervals. A glossary of symbols is included after the Discussion.

**Figure 5.**
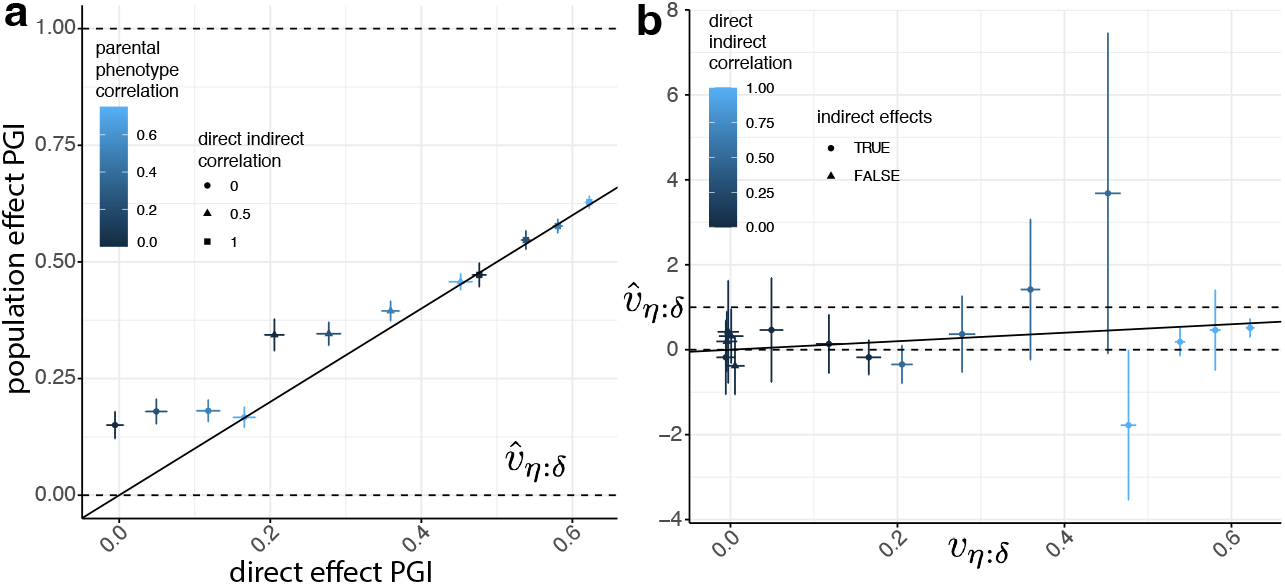
Inferring indirect genetic effects (IGEs) using population effect and noisy direct genetic effect (DGE) PGIs. a) We compare inferences of v_η: δ_ when using a ‘population effect PGI’ (Supplementary Table 6), i.e. a PGI constructed using weights equal to the sum of the DGE and IGE of the SNP plus estimation error, and a DGE PGI, i.e. a PGI constructed using DGEs plus estimation error (Supplementary Table 3). We show results for phenotypes with non-zero IGEs (Supplementary Table 6). Here, the estimation error was set to be equal to the variance of the sum of the true DGEs and IGEs, such that the PGIs captured around one half (or a little less) of the heritability in a random-mating population. b) Across all 16 simulations, estimates of v_η: δ_ when using a DGE PGI that explains only around 1% of the heritability in a random-mating population (Supplementary Table 5). Vertical and horizontal error bars indicate 95% confidence intervals.

While the theory was derived for PGIs constructed from DGEs, most IGE analyses to date have used PGIs constructed from the results of standard GWAS, which estimates ‘population effects’. The population effect of a variant *l* is: *β*_*l*_ = *δ*_*l*_ + *η*_*l*_ + *c*_*l*_, where *c*_*l*_ represents confounding factors that contribute to the population effect of SNP *l*, including from AM^34^. PGIs derived from standard GWAS also have estimation error, *∈*_*βl*_, giving:

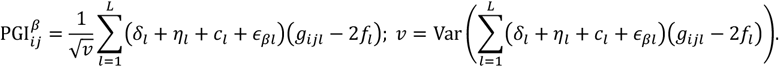

Under a ‘null hypothesis’ of no IGEs or other forms of gene-environment correlation (e.g. population stratification), *β*_*l*_ ≈ *δ*_*l*_/(1 − *r*_*δ*_) (Equation 15 with *k* = 0). This implies the population effects and DGEs are approximately the same up to a scale factor and therefore the true DGE and population effect PGIs are almost perfectly correlated. This implies that PGIs derived from standard GWAS can be used in the above procedure to test the null hypothesis that there are no IGEs or confounding factors other than AM.

To investigate how PGIs constructed from standard GWAS perform in our procedure, we constructed ‘population effect PGIs’ as such:

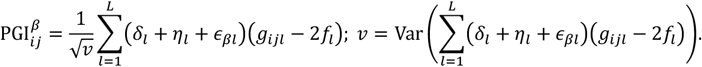

where *∈*_*βl*_ are i.i.d. *N*(0, *v*_*∈β*_) random variables with *v*_*∈β*_ set to multiples of Var_*l*_ (*δ*_*l*_ + *η*_*l*_). (We ignore the confounding due to AM in the population effects since this should be approximately equal to inflating *δ*_*l*_ + *η*_*l*_ by a constant scale factor across variants.)

We compared inference of *v*_*η*: *δ*_ using DGE and population effect PGIs with *k* ≈ 0.5 in Figure 5a (Supplementary Tables 3 and 6). While *v*_*η*: *δ*_ as inferred from a population effect PGI cannot be interpreted as the proportion of phenotypic variance contributed by the IGE component that is correlated with the DGE component, our results show that the population effect PGI can detect the presence of IGEs when the DGE PGI cannot (Figure 5a). This is because there are cases when the DGE and IGE components are uncorrelated or weakly correlated, so a PGI constructed from DGEs does not detect the presence of IGEs. For example, if DGEs and IGEs are uncorrelated and there is no AM, the DGE and IGE components are uncorrelated, and *α*_*δ*_ = 0, so the DGE PGI does not detect IGEs, even though they are present. However, because IGEs contribute to population effects, the IGEs contribute to the weight vector of the population effect PGI, enabling the population effect PGI to detect the presence of IGEs even when DGE and IGE components are uncorrelated (Table 1). A limitation of this analysis is that it ignores the potential impact of population stratification confounding on PGIs derived from standard GWAS.

**Table 2.**
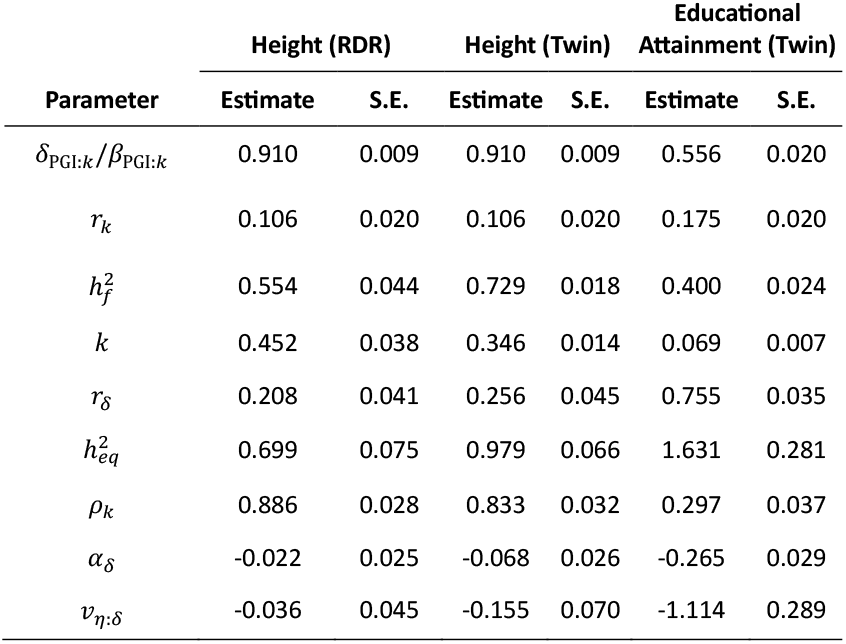
Two generation PGI analysis for height and educational attainment. We applied two-generation PGI analysis (Figure 3) to the results of Okbay et al.^43^ for height and educational attainment (EA) PGIs. Okbay et al. gave estimates of the ratio between direct and population effects of the PGIs, δ_PGI:k_/β_PGI:k_, and of the correlation between parents’ PGIs, r_k_. To complete the inputs to two-generation PGI analysis, we input estimates of 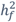 from RDR or from the twin ACE model. We used two different estimates of 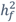 for height: one from a meta-analysis of twin studies^42^, and one from applying RDR to Icelandic data^26^. For EA, we used an estimate of 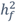 from a meta-analysis of twin studies using the ACE model^53^. By applying a series of non-linear estimating equations (Figure 3 and Methods), we obtained estimates of the fraction of heritability the PGI would explain in a random-mating population, k; the correlation between parents’ direct genetic effect (DGE) components, r_δ_; the equilibrium heritability, 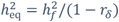; the ratio between direct and population effects of the PGI that would be expected due to assortative mating in the absence of indirect genetic effects (IGEs), ρ_k_; the IGE of the true DGE PGI (that captures all of the heritability), α_δ_; and the contribution to phenotypic variance from by IGE component that is correlated with the DGE, v_η: δ_. A glossary of symbols is included after the Discussion.

### Analysis of PGIs for height and educational attainment

We performed two-generation PGI analysis (Figure 3) for height and EA using results from Okbay et al. 2022 (ref^43^) (Methods and Table 2). Okbay et al. estimated the ratio between PGI direct and population effects and the correlation between parents’ PGIs using PGIs constructed from standard GWAS estimates of ‘population effects’. In addition to the results reported in Okbay et al., we also need an estimate of 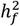. We considered two different estimates of 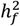 for height: one from a meta-analysis of twin studies using the ACE model^42^, and one from applying RDR to Icelandic data^26^. For educational attainment (EA), we used an estimate from a meta-analysis of twin studies^53^. We did not use an RDR or sib-regression estimate as the only available estimates^23,26^ lacked sufficient precision for reliable inference (Methods and Supplementary Table 7).

Our results using the RDR estimate of 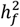 for height show no evidence for IGEs, and the direct to population effect ratio (*δ*_PGI:*k*_/*β*_PGI:*k*_ = 0.910, S.E.=0.009) is statistically indistinguishable from the prediction due to AM alone (without IGEs): *ρ*_*k*_ = 0.886 (S.E.=0.028). We estimated that the correlation between parents’ DGE components is *r*_*δ*_ = 0.208 (S.E.=0.041), around double that for the observed PGI. Using this estimate of *r*_*δ*_, we can adjust the RDR estimate of heritability to give an estimate of the equilibrium heritability of height: 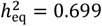 (S.E.=0.075). However, if we use the twin estimate of 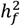, we obtain an implausible result that there are IGEs that are negatively correlated with DGEs. This implausible result could be due to overestimation of heritability by twin studies^26^, which leads to underestimation of *k* and therefore a prediction of the ratio between direct and population effects of the PGI that is too low. An alternative (and not mutually exclusive) explanation is that the estimate of *r*_*k*_ is too high. This could happen when there are confounding factors captured by the height PGI (which is derived from standard GWAS) that are correlated between parents, thereby inflating the correlation between parents’ PGIs beyond what would be expected for a DGE PGI (with no confounding) that explains the same fraction of heritability.

For EA, we estimated that *r*_*δ*_ = 0.755 (S.E.=0.035). This derives from the fact that the estimated correlation between parents’ PGIs was *r*_*k*_ = 0.175 (S.E.=0.020), and the PGI was estimated to explain only around 7% of the heritability in a random-mating population. This very high estimate of *r*_*δ*_ results in an impossibly high estimate of equilibrium heritability (being statistically significantly above 1) and an inference that there are strong IGEs negatively correlated with DGEs. Two plausible explanations for these implausible results are: overestimation of heritability by twin studies^26^, and confounding in the EA GWAS summary statistics^34^ that inflates the correlation between parents’ PGI values. The second phenomenon implies that part of the correlation between parents’ PGIs is due to confounding factors that are correlated between parents — such as social class — in addition to DGEs, leading to a correlation that is higher than would be obtained from a DGE PGI that explains the same fraction of heritability.

## Discussion

In this manuscript, we give a decomposition of the phenotypic variance (Equation 4) under assortative mating (AM) at equilibrium for a phenotype affected by both direct genetic effects (DGEs) and indirect genetic effects (IGEs). We achieve this by a generalization of the method of Crow and Felsenstein^11^ to a phenotype affected by both DGEs and IGEs. The equilibrium variance is expressed in terms of the random-mating variance components as defined by Young et al. in relation to the RDR method of estimating heritability^26^ (Equation 1) and the equilibrium correlations between parents’ DGE and IGE components (Figure 1). We connected our results to estimation of heritability using random variation in realized relatedness between siblings^24^, ‘sib-regression’. We demonstrated theoretically that sib-regression estimates the random-mating genetic variance divided by the equilibrium phenotypic variance, 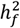, in agreement with Kemper et al.^23^.

An important unsolved problem in human genetics is how to interpret the results from regression models including both proband (phenotyped individual) and parental PGIs^37,39,43^. The expected regression coefficient on the proband PGI, *δ*_PGI_, is called the direct effect of the PGI, and the coefficient on the parental PGI (sum of maternal and paternal PGIs) is called the average non-transmitted coefficient (NTC), *α*_PGI_. We show that, under random mating and for certain special weight vectors, *δ*_PGI_ and *α*_PGI_ are simple functions of the random-mating variance components (Table 1), with *α*_PGI_ reflecting IGEs. When there is AM, the interpretation of the regression coefficients is more complicated. If a PGI would explain only a fraction *k* of the heritability in a random mating population, then *α*_PGI_ captures — in addition to IGEs — the AM-induced correlation of the PGI with the DGE component that the PGI would be uncorrelated with in a random mating population (Equation 17).

We derived theoretical results for PGI analysis under AM at equilibrium. We showed that, when assortative mating is strong and *k* ≪ 1, *α*_PGI_/*δ*_PGI_ can be substantial even in the absence of IGEs (Figure 2). Many studies have estimated how much PGI regression coefficients ‘shrink’ within-family^31,43,44,51^, i.e. how the coefficient on the proband PGI changes when controlling for parental or sibling PGIs. In our framework, this ‘shrinkage’ corresponds to the ratio between direct and population effects, *δ*_PGI_/*β*_PGI_. Our results show that substantial shrinkage can be expected when AM is strong, even in the absence of IGEs (Figure 2). These results argue against naïve interpretations of statistically significant estimates of *α*_PGI_ or substantial ‘shrinkage’ of PGI regression coefficients within-family as demonstrating the influence of IGEs.

In 2018, Kong et al. presented analyses of an educational attainment (EA) PGI in Icelandic data, and estimated that *α*_PGI_/*δ*_PGI_ = 0.427. Kong et al. recognized that *α*_PGI_ could reflect the AM-induced correlation of alleles not transmitted from parents to offspring with transmitted alleles. Kong et al. proposed a technique to adjust for the bias, concluding that the bias was small and therefore that IGEs on EA were substantial. However, the technique proposed by Kong et al. relied upon strong assumptions, including that there had been only one generation of assortative mating and that direct and indirect genetic effects were perfectly correlated. Furthermore, Kong et al. did not account for uncertainty in input parameters or in their adjustment.

We develop a procedure (Figure 3 and Methods), implemented in *snipar*, that accounts for AM by combining the results of two-generation PGI analysis with a family-based heritability estimate 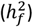 and an estimate of the correlation between parents’ PGIs (*r*_*k*_). When applied to PGIs constructed from DGE estimates, this produces estimates of the correlation between parents’ DGE components, *r*_*δ*_, the equilibrium heritability, 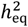, and the contribution to phenotypic variance from the IGE component that is correlated with the DGE component, *v*_*η*: *δ*_. Simulations show that this gives approximately unbiased estimates along with accurate standard errors when all the input parameters are estimated precisely (Methods and Supplementary Tables 2-7). Unlike the procedure developed by Kong et al., our procedure does not assume perfectly correlated DGEs and IGEs.

The theory underlying the method is derived for a PGI constructed using DGEs as weights. However, almost all investigations of IGEs have been performed using PGIs constructed using ‘population effect’ estimates from standard GWAS^31,43,51^. The advantage of such PGIs over PGIs constructed from DGE estimates from family-based GWAS is greater statistical power due to larger sample sizes and the fact that ‘population effects’ capture both DGEs and IGEs. Under a ‘null model’ of AM but no IGEs, DGEs and population effects of variants should be approximately perfectly correlated, implying that population effect PGIs can be used in our procedure (Figure 3) to test a null hypothesis that there are no IGEs (or other confounding factors beyond AM). Simulations indicate that population effect PGIs can detect the influence of IGEs in situations when DGE PGIs do not (Figure 4), since they will do so even when DGE and IGE components are uncorrelated.

We applied our method to results on EA and height population effect PGIs from Okbay et al.^43^ (Table 2). The height PGI results — using an estimate of 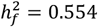 from applying RDR to Icelandic data^54^ — indicated no evidence for IGEs on height and gave an estimate of the equilibrium heritability of height of 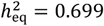 (S.E. = 0.075). Our estimation procedure produced nonsensical results when applied to the EA PGI, including that the equilibrium heritability is above 1. The equilibrium heritability estimate is so high because our estimate of the correlation between parents’ DGE components is so high: *r*_*δ*_ = 0.755 (S.E.=0.035). This derives from the fact that the correlation between the parents’ observed PGIs is high, *r*_*k*_ = 0.175 (S.E.=0.020), even though the PGI is estimated to explain only around 7% of the heritability in a random-mating population.

Our procedure for estimating the correlation between parents’ underlying DGE components, *r*_*δ*_, accounts for PGI-environment correlation — such as from IGEs and population stratification — although not for confounding in the GWAS summary statistics used to construct the PGI. This is an advantage over a recently proposed structural equation model, *rgensi*^13^, which uses siblings and their in-laws in combination with a PGI to estimate *r*_*δ*_. Our estimate of *r*_*δ*_ for EA was higher than from *rgensi* applied to an EA PGI in the Norwegian Mother, Father and Child Cohort Study (MoBa), which estimated *r*_*δ*_ = 0.37 (95% confidence interval: [0.21,0.67]). The primary reason for this difference is that the *rgensi* model does not estimate the effect of the EA PGI within-family — which is how our method accounts for PGI-environment correlation — and thus obtained a much higher estimate for the fraction of heritability explained by the PGI: 25% (95% confidence interval: [12%,42%]).

While the simulation results indicated that using a population effect PGI in our procedure produces reasonably accurate results, the simulations did not consider the impact of population stratification confounding in population effect estimates. We hypothesize that confounding factors in the EA GWAS have inflated the correlation between parents’ EA PGIs beyond that which would be expected for a DGE PGI (without confounding) that explains the same amount of heritability. This would have the effect of overestimating the correlation between parents’ DGE components and therefore the equilibrium heritability.

This manuscript has focused on estimating a consequence of cultural/vertical transmission from parents to offspring, the IGEs that are induced when the parental phenotypes affecting the offspring through the environment are themselves heritable. Previous research on estimation of cultural/vertical transmission includes the cultural transmission model of Cloninger, Rice, and Reich^18^, extended twin and family designs (ETDFs) such as the ‘stealth’ and ‘cascade’ models, and the structural equation model of Balbona et al.^33^. Unlike previous research, our results do not rely upon a particular model of assortative mating. A difference from previous models is that we do not model the total contribution of vertical/cultural transmission from parents to offspring, only the heritable component of the parental phenotype that influences offspring through the environment. While modelling the total contribution of vertical/cultural transmission is important for predicting phenotypic correlations between relatives, it is less important for PGI analyses and estimation of IGEs.

Our method does not make any assumptions about the parental phenotype through which cultural/vertical transmission from parents to offspring operates and can model arbitrary correlations between DGEs and IGEs. This is in contrast to most previous research — except for the model from Cloninger, Rice, and Reich, which, while flexible, cannot separately identify genetic and cultural transmission when fit to phenotypic correlations between relatives^19^. ETFDs can separately identify genetic and cultural transmission, but these models generally assume vertical/cultural transmission operates through an observed phenotype. The same is true of the structural equation model of Balbona et al., which is the only other method that uses PGIs.

This manuscript provides a general theoretical framework for understanding the joint impact of AM and IGEs on phenotypic variance components, heritability estimation, and PGI analyses. As more genetic data on families becomes available, we will obtain more powerful family-based GWAS summary statistics, enabling us to apply our estimation procedure to DGE PGIs. We will also obtain more precise heritability estimates from sib-regression^24^ and RDR^54^. Together, these will enable application of the theoretical results and methodology developed here to quantify the impact of IGEs and AM on phenotype variation.

## Glossary

|  |  |
| --- | --- |
| $\delta_l$ | Direct genetic effect (DGE) of variant $l$ |
| $\eta_l$ | Average parental indirect genetic effect (IGE) of variant $l$ |
| $g_{ijl}$ | Genotype of sibling $j$ in family $i$ at variant $l$. Variants are assumed to be bi-allelic, with genotypes counting the number of copies of the allele (0, 1, or 2) with frequency $f_l$ |
| $g_{p(i)l}$ | Genotype of the father in family $i$ at variant $l$ |
| $g_{m(i)l}$ | Genotype of the mother in family $i$ at variant $l$ |
| $g_{\text{par}(i)l}$ | Combined parental genotype for family $i$ at variant $l$: $g_{\text{par}(i)l} = g_{p(i)l} + g_{m(i)l}$ |
| $Y_{ij}$ | Phenotype of sibling $j$ in family $i$. |
| $\Delta_{ij}, \Delta_{p(i)}, \Delta_{m(i)}$ | Direct genetic effect (DGE) component of sibling $j$ in family $i$, the father in family $i$, and the mother in family $i$:<br>$\Delta_{ij} = \sum_{l=1}^L \delta_l (g_{ijl} - 2f_l); \Delta_{p(i)} = \sum_{l=1}^L \delta_l (g_{p(i)l} - 2f_l);$ $\Delta_{m(i)} = \sum_{l=1}^L \delta_l (g_{m(i)l} - 2f_l).$ |
| $\Delta_{\text{par}(i)}$ | The combined parental DGE component:<br>$\Delta_{\text{par}(i)} = \Delta_{p(i)} + \Delta_{m(i)}.$ |
| $\eta_{p(i)}, \eta_{m(i)}$ | Paternal and maternal indirect genetic effect (IGE) components:<br>$\eta_{p(i)} = \sum_{l=1}^L \eta_l (g_{p(i)l} - 2f_l), \text{ and } \eta_{m(i)} = \sum_{l=1}^L \eta_l (g_{m(i)l} - 2f_l).$ |
| $r_\delta$ | Correlation between parents' DGE components:<br>$r_\delta = \text{Corr}(\Delta_{p(i)}, \Delta_{m(i)}).$ |
| $r_\eta$ | Correlation between parents' IGE components:<br>$r_\eta = \text{Corr}(\eta_{p(i)}, \eta_{m(i)}).$ |
| $r_{\delta\eta}^c$ | Within-parent correlation between DGE and IGE components:<br>$r_{\delta\eta}^c = \text{Corr}(\eta_{p(i)}, \delta_{p(i)}) = \text{Corr}(\eta_{m(i)}, \delta_{m(i)}).$ |
| $r_{\delta\eta}^\tau$ | Between-parent correlation between DGE and IGE components:<br>$r_{\delta\eta}^\tau = \text{Corr}(\eta_{p(i)}, \delta_{m(i)}) = \text{Corr}(\eta_{m(i)}, \delta_{p(i)}).$ |
| $v_g$ | Random-mating variance of DGE component:<br>$v_g = \text{Var}(\Delta_{ij}) = 2 \sum_{l=1}^L \delta_l^2 f_l (1 - f_l).$ |
| $v_{e\sim g}$ | Random-mating variance of parental IGE component:<br>$v_{e\sim g} = \text{Var}(\eta_{p(i)} + \eta_{m(i)}) = 4 \sum_{l=1}^L \eta_l^2 f_l (1 - f_l).$ |
| $c_{ge}$ | Random-mating variance due to covariance between DGE and IGE components:<br>$c_{ge} = 2\text{Cov}(\delta_{ij}, \eta_{p(i)} + \eta_{m(i)}) = 4 \sum_{l=1}^L \delta_l \eta_l f_l (1 - f_l).$ |
| $v_y^{\text{eq}}$ | Equilibrium phenotypic variance. |
| $v_g^{\text{eq}}$ | Equilibrium variance of DGE component: $v_g^{\text{eq}} = \frac{v_g}{1-r_\delta}$. |
| $h_{\text{eq}}^2$ | Equilibrium heritability: $h_{\text{eq}}^2 = v_g^{\text{eq}} / v_y^{\text{eq}}$ |
| $v_{e\sim g}^{\text{eq}}$ | Equilibrium variance of parental IGE component: $v_{e\sim g}^{\text{eq}} = \frac{1+r_\eta}{1-r_\eta} v_{e\sim g}$ |
| $c_{ge}^{\text{eq}}$ | Equilibrium variance due to covariance between DGE and IGE components:<br>$c_{ge}^{\text{eq}} = (r_{\delta\eta}^c + r_{\delta\eta}^\tau) \sqrt{\frac{2v_{e\sim g} v_g}{(1-r_\eta)(1-r_\delta)}}.$ |
| $r_{\delta\eta}^0$ | Random-mating correlation between DGE and IGE components. |
| $R_{ijk}$ | Realized relatedness between siblings $j$ and $k$ in family $i$. |
| $h_f^2$ | Within-family heritability estimand at equilibrium (as from MZ-DZ twin comparisons, relatedness disequilibrium regression, and sib-regression):<br>$h_f^2 = v_g/v_y^{\text{eq}} = (1 - r_\delta)h_{\text{eq}}^2$ |
| $\text{PGI}_{ij}$ | Polygenic index of sibling $j$ in family $i$ |
| $\text{PGI}_{\text{par}(i)}$ | Combined parental polygenic index in family $i$:<br>$\text{PGI}_{\text{par}(i)} = \text{PGI}_{p(i)} + \text{PGI}_{m(i)}$, where $\text{PGI}_{p(i)}$ and $\text{PGI}_{m(i)}$ are, respectively, the paternal and maternal PGIs |
| $\delta_{\text{PGI}}, \alpha_{\text{PGI}}$ | Direct effect and average non-transmitted coefficient (NTC) of a PGI defined by the regression: $Y_{ij} = \delta_{\text{PGI}}\text{PGI}_{ij} + \alpha_{\text{PGI}}\text{PGI}_{\text{par}(i)} + \epsilon_{ij}$. |
| $\beta_{\text{PGI}}$ | Population effect of PGI defined by the regression: $Y_{ij} = \beta_{\text{PGI}}\text{PGI}_{ij} + \epsilon_{ij}$. |
| $\alpha_\delta$ | Average non-transmitted coefficient of the true direct genetic effect PGI, i.e. of the DGE component, defined by the regression:<br>$Y_{ij} = \Delta_{ij} + \alpha_\delta\Delta_{\text{par}(i)} + \epsilon_{ij}$ |
| $v_{\eta:\delta}$ | The contribution to phenotypic variance from the IGE component correlated with the DGE component (both its variance and variance due to covariance with the DGE component):<br>$v_{\eta:\delta} = 2(1 + r_\delta)\alpha_\delta(1 + \alpha_\delta)h_{\text{eq}}^2$ |
| $\text{PGI}_{ij}^{\delta_k}$ | Variance-standardized direct effect PGI that explains a fraction $k$ of the heritability in a random-mating population — for example, with $\delta_l = \delta$ and $f_l = f$ for all $l$,<br>$\text{PGI}_{ij}^{\delta_k} = \frac{\delta}{\sqrt{v_k}} \sum_{l=1}^{kL} (g_{ijl} - 2f), \quad v_k = \text{Var}\left(\sum_{l=1}^{kL} (g_{ijl} - 2f)\right).$ |
| $\delta_{\text{PGI}:k}, \alpha_{\text{PGI}:k}$ | The direct effect and average non-transmitted coefficient of a DGE PGI that explains a fraction $k$ of the heritability in a random-mating population, defined by a regression with phenotype and PGI standardized to have variance 1:<br>$\frac{Y_{ij}}{\sqrt{v_y^{\text{eq}}}} = \delta_{\text{PGI}:k}\text{PGI}_{ij}^{\delta_k} + \alpha_{\text{PGI}:k}\text{PGI}_{\text{par}(i)}^{\delta_k} + \epsilon_{ij}.$ |
| $\beta_{\text{PGI}:k}$ | The population effect of a DGE PGI that explains a fraction $k$ of the heritability in a random-mating population, defined by a regression with phenotype and PGI standardized to have variance 1:<br>$\frac{Y_{ij}}{\sqrt{v_y^{\text{eq}}}} = \beta_{\text{PGI}:k}\text{PGI}_{ij}^{\delta_k} + \epsilon_{ij}.$ |
| $r_k$ | The correlation between parents' DGE PGIs that explain a fraction $k$ of the heritability in a random-mating population:<br>$r_k = \text{Corr}\left(\text{PGI}_{p(i)}^{\delta_k}, \text{PGI}_{m(i)}^{\delta_k}\right).$ |
| $\rho_k$ | The ratio between direct and population effects of a DGE PGI that explains a fraction $k$ of the heritability in a random mating population in a model without indirect genetic effects: $\rho_k = \frac{\delta_{\text{PGI}:k}}{\beta_{\text{PGI}:k}} = 1 - (1 - k)r_\delta$. |

## Data Availability

Simulated populations (genotype and phenotype data) will be made available for download on publication of the final version of this manuscript.

## Code Availability

The code for performing simulations of phenotypes affected by DGEs, parental IGEs, and AM is available as a command line script (*simulate.py*) in *snipar* (https://github.com/AlexTISYoung/snipar). The code for performing two-generation PGI analysis accounting for AM (Figure 3) is available as a command line script (*pgs.py*) in *snipar*. We provide a tutorial on simulating data and performing two-generation PGI analysis here: https://snipar.readthedocs.io/en/latest/simulation.html

The code for the specific simulations and PGI analyses described in this paper is available here: https://github.com/AlexTISYoung/snipar/tree/simulate.

## Acknowledgments

The study was supported by Open Philanthropy (010623-00001 and 2019-198171) and the National Institute on Aging/National Institutes of Health through grants R24-AG065184 and R01-AG042568 (to the University of California, Los Angeles). I thank Daniel J. Benjamin, Patrick Turley, James J. Lee, Greg Clark, Peter Visscher, Richard Border, and Dalton Conley for helpful comments and suggestions.

## Methods

### Estimating the fraction of heritability a PGI would explain in a random-mating population

In real-world applications, we do not know *k*, the fraction of heritability the PGI would explain in a random-mating population. In Supplementary Note Section 8, we show that

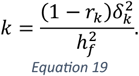

To estimate *k*, consider that we have unbiased and statistically independent estimates of *r*_*k*_, given by 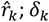; *δ*_*k*_, given by 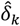; and 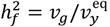, given by 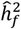. The estimate of 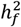 could come from sib-regression, RDR, or from the ACE twin model. In Supplementary Note Section 8, we propose a bias-corrected estimator of *k*,

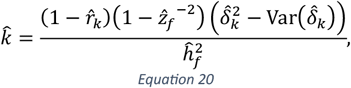

where 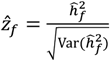.

### Estimating the correlation between parents’ DGE components and equilibrium heritability

In order to make inferences about the impact of assortative mating on the DGE component, we need to estimate

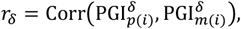

the correlation between the parents’ DGE components. We estimate *r*_*k*_ by (Supplementary Note Section 7.2):

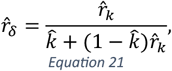

with 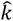 as defined above. Provided that the estimators of *k* and *r*_*k*_ are consistent, this estimator of *r*_*δ*_ is consistent. However, the estimator is biased, and becomes unstable when the estimates of *k* and *r*_*k*_ are noisy, or when the denominator *k* + (1 − *k*)*r*_*k*_ is close to zero (Supplementary Table 7).

As shown above, 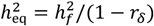, which implies one can obtain an estimate of 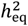 by combining an estimate of 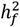 with an estimate of *r*_*δ*_:

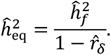

### Estimating the ratio between direct and population effects due to AM

To estimate what the ratio between direct and population effects of a PGIs would be under a model with AM at equilibrium but without IGEs, one can combine *k* and *r*_*δ*_ as estimated above (Equations 20 and 21):

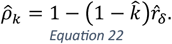

### Parameter estimate bias and sampling error estimation

Since the parameters are estimated using a series of non-linear equations, we expect the estimators to be biased even when the input parameter estimates are themselves unbiased. Furthermore, standard errors are approximated using the Delta method (Supplementary Note Section 9), which can be inaccurate when the input parameters are noisy. We simulated input parameter data emulating that which could be obtained from performing two-generation PGI analysis using either 10,000 or 50,000 independent trios along with a statistically independent and unbiased estimate of 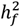 (Supplementary Note Section 10). We found that parameter estimates were approximately unbiased and standard errors were accurate when all input parameters were estimated precisely (Supplementary Table 7), i.e. the ratio between the true parameter value and its standard error was greater than around 3. When, for example, the estimate of 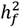 was noisy and/or *k* was small, parameter estimates could be substantially biased and standard errors inaccurate.

### Simulation of populations and phenotypes

We simulated 16 phenotypes with varying parameters using the *simulate.py* module in *snipar* (Supplementary Table 1). Each phenotype was simulated by first simulating 60,000 independent individuals at 1,000 causal SNPs. SNP minor-allele frequencies (MAFs), *f*_*l*_, were simulated from a distribution with density proportional to 1/*f*_*l*_, ranging from *f*_*l*_ = 0.05 to *f*_*l*_ = 0.5. We chose this distribution since MAFs are expected to have a distribution proportional to 1/*f* for a randommating population of constant effective size^55^. For each SNP *l*, the genotype of each individual was drawn independently from a Binomial(2, *f*_*l*_) distribution. The 60,000 individuals were then randomly paired into 30,000 parent-pairs. For each parent pair, we simulated meiosis, with independent segregation between SNPs, to produce one male and one female (full-sibling) offspring. To simulate the DGE and IGE components in this first offspring generation produced by random-mating, we drew DGEs and IGEs of individual SNPs independently from a bivariate normal distribution:

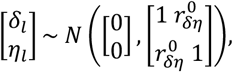

where 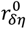 was set to 0, 0.5, or 1 depending upon the simulation. The DGE and IGE components were then scaled to have the desired variance:

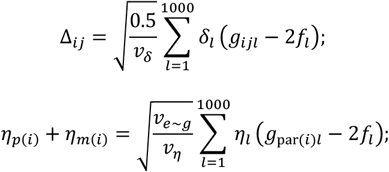

where

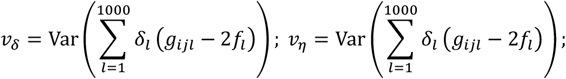

and *v*_*e*∼*g*_ was set to 0.125 or 0 depending on the simulation. The phenotype of sibling *j* in family *i* was constructed as:

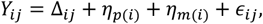

where *∈*_*ij*_ were simulated as independent random variables with 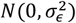 distribution. The variance of the residuals, 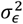, was set such that the overall phenotypic variance in the first generation produced by random-mating was 1:

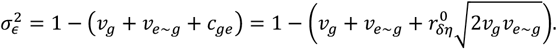

This implies that the heritability in the random-mating population was 0.5 for all phenotypes. For phenotypes with IGEs, the proportion of variance explained by parental IGEs was 0.125, and the proportion of variance explained by covariance between DGEs and IGEs was 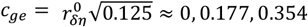 for 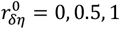.

Subsequent generations were produced by random-mating or AM. AM was simulated by matching males and females into parent-pairs according to their rank on a noisy observation of their phenotype value:

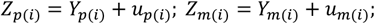

where *u*_*p*(*i*)_ and *u*_*m*(*i*)_ were simulated as independent random variables with distribution 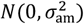. The noise level 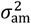 was set such that Corr(*Y*_*p*(*i*),_ *Y*_*m*(*i*)_) ≈ *r*_*y*_ in each generation, which is achieved when 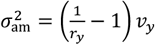, where *v*_*y*_ is the phenotypic variance in the parental generation. For each set of IGE parameters, we performed simulations with different strengths of AM: *r*_*y*_ = 0, 0.25, 0.5, 0.75. We performed 20 generations of random (*r*_*y*_ = 0) or assortative mating (*r*_*y*_ > 0), recording the phenotype values and true variance components in each generation. We retained the genotypes of the final two generations in order to perform PGI analyses.

## References

1. Galton, F. Regression Towards Mediocrity in Hereditary Stature. J. Anthropol. Inst. G. B. Irel. 15, 246–263 (1886).

2. Fisher, R. A. The Correlation between Relatives on the Supposition of Mendelian Inheritance. Trans. R. Soc. Edinb. 52, 399–433 (1918).

3. Provine, W. B. The Origins of Theoretical Population Genetics: With a New Afterword. (University of Chicago Press, 2001).

4. Lynch, M. & Walsh, B. Genetics and Analysis of Quantitative Traits. Genetics and Analysis of Quantitative Traits (Sinauer Associates, Inc., 1998).

5. Young, A. I. & Durbin, R. Estimation of Epistatic Variance Components and Heritability in Founder Populations and Crosses. Genetics 198, 1405–1416 (2014).

6. Falconer, D. S. & Mackay, T. F. C. Introduction to Quantitative Genetics. (Longman, 1996).

7. Crow, J. F. & Kimura, M. An introduction to population genetics theory. The Blackburn Press (New York, Evanston and London: Harper & Row, Publishers, 1970).

8. Nagylaki, T. Assortative mating for a quantitative character. J. Math. Biol. 16, 57–74 (1982).

9. Sunde, H. F. et al. Genetic similarity between relatives provides evidence on the presence and history of assortative mating. bioRxiv (2023) doi:10.1101/2023.06.27.546663.

10. Clark, G. The inheritance of social status: England, 1600 to 2022. Proc. Natl. Acad. Sci. 120, e2300926120 (2023).

11. Crow, J. F. & Felsenstein, J. The effect of assortative mating on the genetic composition of a population. Eugen. Q. 15, 85–97 (1968).

12. Keller, M. C. et al. Modeling Extended Twin Family Data I: Description of the Cascade Model. Twin Res. Hum. Genet. 12, 8–18 (2009).

13. Torvik, F. A. et al. Modeling assortative mating and genetic similarities between partners, siblings, and in-laws. Nat. Commun. 13, 1108 (2022).

14. Cavalli-Sforza, L. L. & Feldman, M. W. Cultural versus biological inheritance: Phenotypic transmission from parents to children (A theory of the effect of parental phenotypes on children’s phenotypes). Am. J. Hum. Genet. 25, 618–637 (1973).

15. Cavalli-Sforza, L. L. & Feldman, M. W. The evolution of continuous variation. III. Joint transmission of genotype, phenotype and environment. Genetics 90, 391–425 (1978).

16. Feldman, M. W. & Laland, K. N. Gene-culture coevolutionary theory. Trends Ecol. Evol. 11, 453–457 (1996).

17. Rice, J., Cloninger, C. R. & Reich, T. Multifactorial inheritance with cultural transmission and assortative mating. I. Description and basic properties of the unitary models. Am. J. Hum. Genet. 30, 618–643 (1978).

18. Cloninger, C. R., Rice, J. & Reich, T. Multifactorial inheritance with cultural transmission and assortative mating. II. a general model of combined polygenic and cultural inheritance. Am. J. Hum. Genet. 31, 176–98 (1979).

19. Ruby, J. G. et al. Estimates of the Heritability of Human Longevity Are Substantially Inflated due to Assortative Mating. Genetics 210, 1109–1124 (2018).

20. Collado, M. D., Ortuño-Ortín, I. & Stuhler, J. Estimating Intergenerational and Assortative Processes in Extended Family Data. Rev. Econ. Stud. 90, 1195–1227 (2023).

21. Truett, K. R. et al. A model system for analysis of family resemblance in extended kinships of twins. Behav. Genet. 24, 35–49 (1994).

22. Keller, M. C., Medland, S. E. & Duncan, L. E. Are extended twin family designs worth the trouble? A comparison of the bias, precision, and accuracy of parameters estimated in four twin family models. Behav. Genet. 40, 377–393 (2010).

23. Kemper, K. E. et al. Phenotypic covariance across the entire spectrum of relatedness for 86 billion pairs of individuals. Nat. Commun. 12, 1050 (2021).

24. Visscher, P. M. et al. Assumption-free estimation of heritability from genome-wide identity-by-descent sharing between full siblings. PLoS Genet. 2, e41 (2006).

25. Yang, J. et al. Common SNPs explain a large proportion of the heritability for human height. Nat. Genet. 42, 565–9 (2010).

26. Young, A. I. et al. Relatedness disequilibrium regression estimates heritability without environmental bias. Nat. Genet. 50, 1304–1310 (2018).

27. Zaitlen, N. et al. Using Extended Genealogy to Estimate Components of Heritability for 23 Quantitative and Dichotomous Traits. PLoS Genet. 9, e1003520 (2013).

28. Yengo, L. et al. Imprint of assortative mating on the human genome. Nat. Hum. Behav. 2, 948–954 (2018).

29. Conley, D. et al. Assortative mating and differential fertility by phenotype and genotype across the 20th century. Proc. Natl. Acad. Sci. 113, 6647–6652 (2016).

30. Robinson, M. R. et al. Genetic evidence of assortative mating in humans. Nat. Hum. Behav. 1, 0016 (2017).

31. Kong, A. et al. The nature of nurture: Effects of parental genotypes. Science 359, 424– 428 (2018).

32. Wang, B. et al. Robust genetic nurture effects on education: A systematic review and meta-analysis based on 38,654 families across 8 cohorts. Am. J. Hum. Genet. 108, 1780–1791 (2021).

33. Balbona, J. V., Kim, Y. & Keller, M. C. Estimation of Parental Effects Using Polygenic Scores. Behav. Genet. (2021) doi:10.1007/s10519-020-10032-w.

34. Young, A. I. et al. Mendelian imputation of parental genotypes improves estimates of direct genetic effects. Nat. Genet. 54, 897–905 (2022).

35. Young, A. I., Benonisdottir, S., Przeworski, M. & Kong, A. Deconstructing the sources of genotype-phenotype associations in humans. Science 365, 1396–1400 (2019).

36. Carey, G. Sibling imitation and contrast effects. Behav. Genet. 16, 319–341 (1986).

37. Nivard, M. et al. Neither nature nor nurture: Using extended pedigree data to elucidate the origins of indirect genetic effects on offspring educational outcomes. BioRxiv (2022).

38. Shen, H. & Feldman, M. W. Genetic nurturing, missing heritability, and causal analysis in genetic statistics. Proc. Natl. Acad. Sci. 117, 25646–25654 (2020).

39. Veller, C. & Coop, G. Interpreting population and family-based genome-wide association studies in the presence of confounding. BioRxiv 2023.02.26.530052 (2023) doi:10.1101/2023.02.26.530052.

40. Border, R. et al. Assortative mating biases marker-based heritability estimators. Nat. Commun. 13, 1–10 (2022).

41. Boomsma, D., Busjahn, A. & Peltonen, L. Classical twin studies and beyond. Nat. Rev. Genet. 3, 872–82 (2002).

42. Polderman, T. J. C. et al. Meta-analysis of the heritability of human traits based on fiNy years of twin studies. Nat. Genet. 47, 702–709 (2015).

43. Okbay, A. et al. Polygenic prediction of educational attainment within and between families from genome-wide association analyses in 3 million individuals. Nat. Genet. 54, 437– 439 (2022).

44. Howe, L. J. et al. Within-sibship genome-wide association analyses decrease bias in estimates of direct genetic effects. Nat. Genet. 54, 581–592 (2022).

45. Becker, J. et al. Resource profile and user guide of the Polygenic Index Repository. Nat. Hum. Behav. 5, 1744–1758 (2021).

46. Mostafavi, H. et al. Variable prediction accuracy of polygenic scores within an ancestry group. Elife 30, e48376 (2020).

47. Martin, A. R. et al. Clinical use of current polygenic risk scores may exacerbate health disparities. Nat. Genet. 2019 514 51, 584–591 (2019).

48. Young, A. I. Discovering missing heritability in whole-genome sequencing data. Nat. Genet. 54, 224–226 (2022).

49. Young, A. I. Solving the missing heritability problem. PLoS Genet. 15, 1–7 (2019).

50. Wainschtein, P., Jain, D., Zheng, Z. & Working, T. A. Assessing the contribution of rare variants to complex trait heritability from whole genome sequence data. Nat. Genet. (2022).

51. Selzam, S. et al. Comparing Within- and Between-Family Polygenic Score Prediction. Am. J. Hum. Genet. 105, 351 (2019).

52. Lee, J. J. et al. Gene discovery and polygenic prediction from a genome-wide association study of educational attainment in 1.1 million individuals. Nat. Genet. 50, 1112–1121 (2018).

53. Branigan, A. R., Mccallum, K. J. & Freese, J. Variation in the heritability of educational attainment: An international meta-analysis. Soc. Forces 92, 109–140 (2013).

54. Young, A. Relatedness disequilibrium regression estimates heritability without environmental bias Origins of heritability estimation. (2018).

55. Uricchio, L. H. Evolutionary perspectives on polygenic selection, missing heritability, and GWAS. Hum. Genet. 139, 5–21 (2020).

