## Supplementary Figures and Note for "Estimation of indirect genetic effects and heritability under assortative mating"

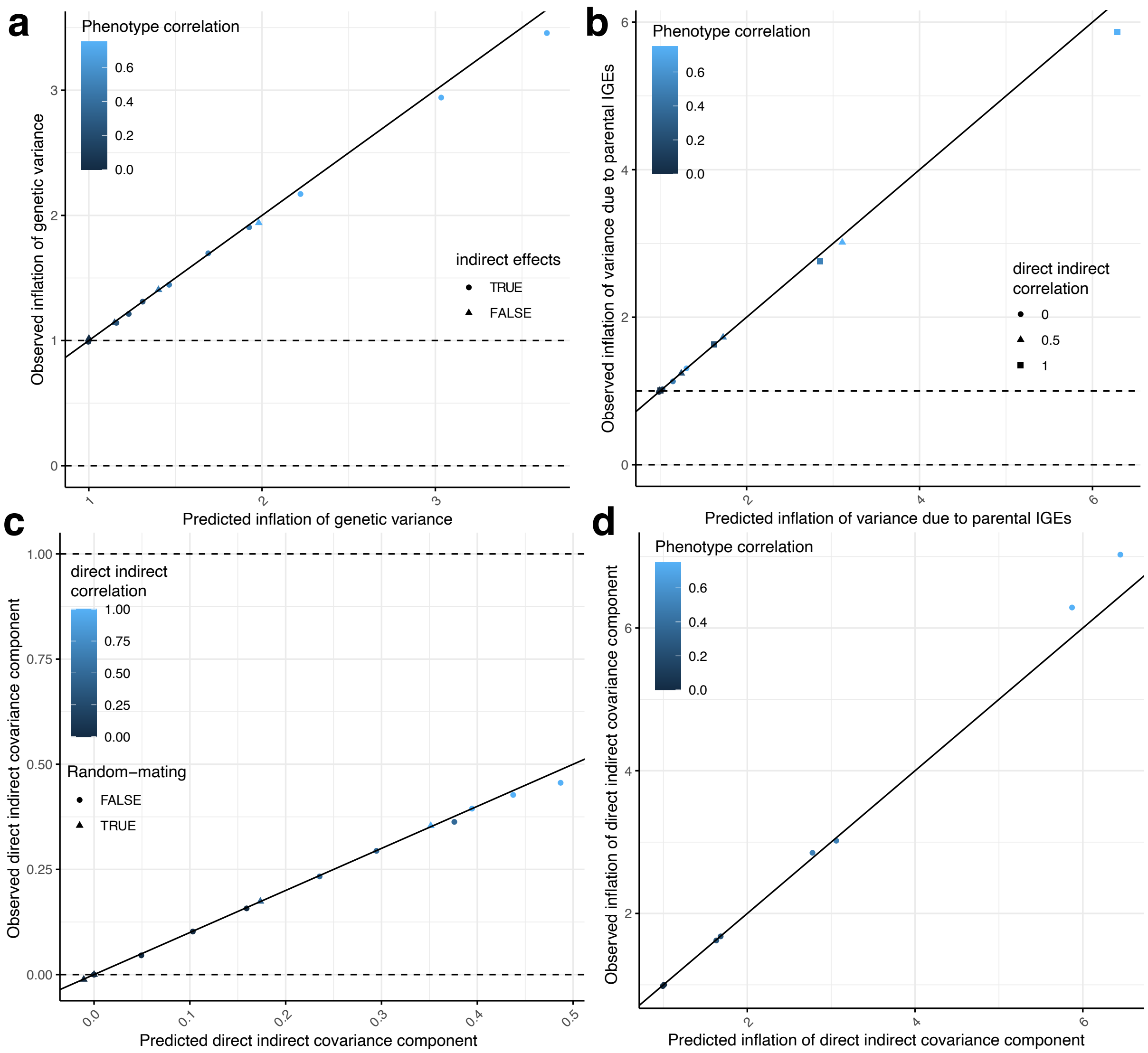


***Supplementary Figure 1. Equilibrium phenotypic variance components: theory vs. observations.*** *We simulated 16 phenotypes with differing indirect genetic effect (IGE) parameters and strengths of assortative mating (Methods). a) We compare the predicted inflation of genetic variance due to assortative mating at equilibrium, $1/(1-r_\delta)$, to the observed inflation, i.e. the genetic variance after 20 generations of mating compared to the genetic variance in the first generation produced by random mating. b) Comparisons of predicted inflation of variance due to parental IGEs, $(1+r_\eta)/(1-r_\eta)$, to the observed inflation after 20 generations of mating. c) We compare the predicted phenotypic variance component due to covariance between direct genetic effect (DGE) and IGE components, $\left(r^c_{\delta\eta}+r^\tau_{\delta\eta}\right)\sqrt{\frac{2v_{e\sim g}v_g}{(1-r_\eta)(1-r_\delta)}}$, to the observed variance component. d) For phenotypes with non-zero correlation DGEs and IGEs, we compare the predicted inflation of the variance component due to covariance between DGE and IGE components, $\frac{r^c_{\delta\eta}+r^\tau_{\delta\eta}}{r^c_{\delta\eta}-r^\tau_{\delta\eta}}$, to the observed inflation of this variance component after 20 generations of mating.*

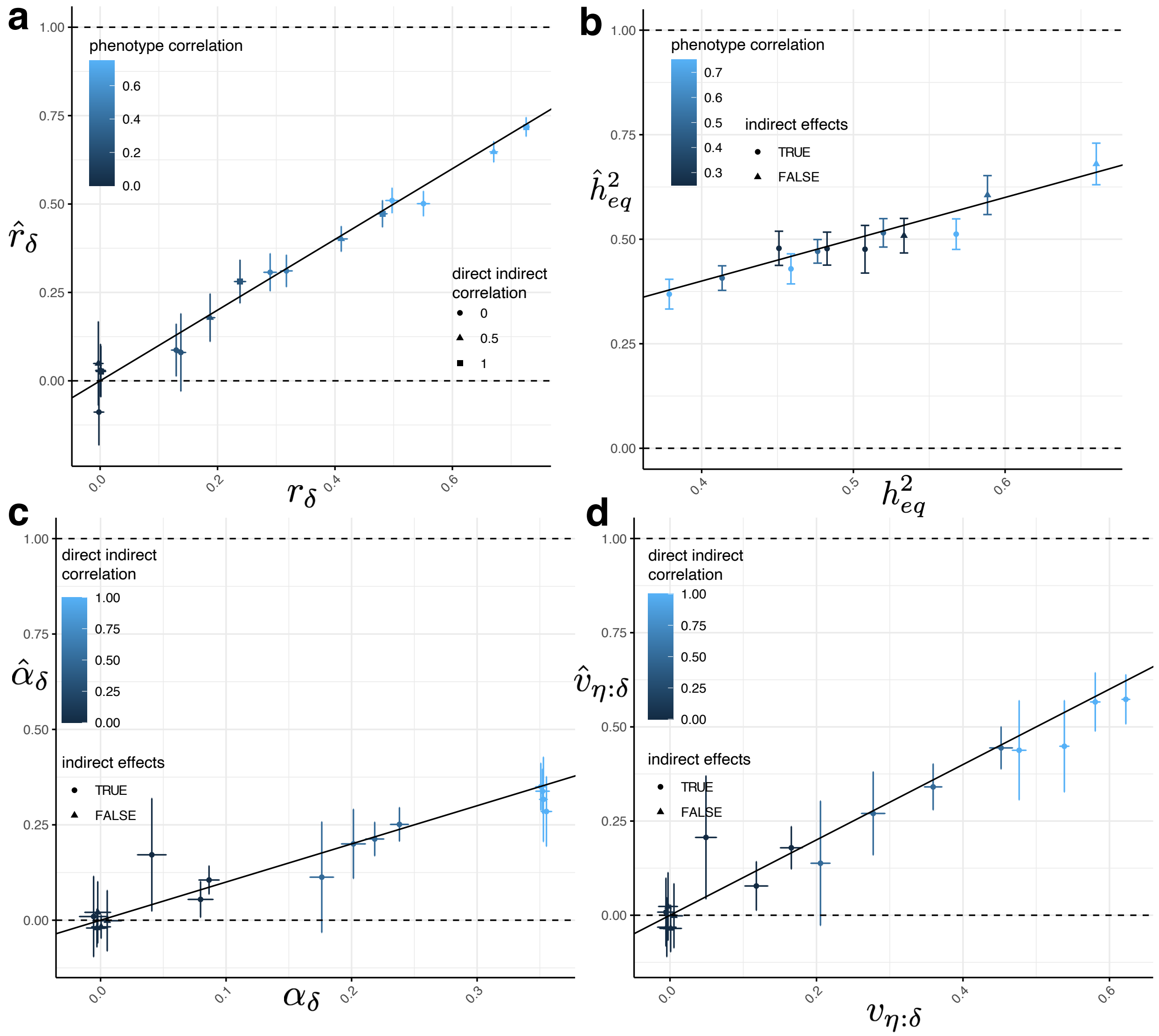


***Supplementary Figure 2. Simulation results for a direct genetic effect (DGE) PGI with noise level 10.*** *Across 16 simulated phenotypes (Methods), we computed PGIs using weights equal to the true direct genetic effects plus a noise term equal to 10x the variance to the true DGEs. This resulted in a DGE PGI that explained approximately 9% of the heritability in a random-mating population (Supplementary Table 4). We performed two-generation PGI analysis (Methods and Figure 3) in order to estimate a)* $r_\delta$*, the correlation between parents' true DGE components (that explain all the heritability); b)* $h^2_{eq}$*, the equilibrium heritability; c)* $\alpha_\delta$*, the indirect genetic effect of the true DGE PGI; and d) the proportion of phenotypic variance contributed by the IGE component that is correlated with the DGE component,* $v_{\eta:\delta}$*. Vertical and horizontal error bars indicate 95% confidence intervals.*

### Supplementary Note for: Estimation of indirect genetic effects and heritability under assortative mating

#### 1 Phenotype model

We consider a model with direct genetic effects (DGEs) and indirect genetic effects (IGEs) from parents:

$$Y_{ij} = \Delta_{ij} + \eta_{p(i)} + \eta_{m(i)} + \epsilon_{ij}$$

where $Y_{ij}$ is the phenotype of individual $i$ in family $j$; $\epsilon_{ij}$ is the residual that is uncorrelated with $\Delta_{ij}, \eta_{p(i)}, \eta_{m(i)}$ and has variance $\sigma_\epsilon^2$;

$$\Delta_{ij} = \sum_{l=1}^{L} \delta_l \left(g_{ijl} - 2f_l\right); \eta_{p(i)} = \sum_{l=1}^{L} \eta_l \left(g_{p(i)l} - 2f_l\right); \eta_{m(i)} = \sum_{l=1}^{L} \eta_l \left(g_{m(i)l} - 2f_l\right);$$

where $g_{ijl}$ is the genotype of individual $j$ in family $i$ at locus $l$; $g_{p(i)l}$ is the genotype of the father in family $i$ at locus $l$; $g_{m(i)l}$ is the genotype of the mother in family $i$ at locus $l$; and $f_l$ is the allele frequency at locus $l$ such that $\mathbb{E}\left[g_{ijl}\right] = 2f_l$ (assumed to be constant across generations); the DGE of locus $l$ is given by $\delta_l$; and the paternal and maternal IGEs at locus $l$ are given by $\eta_l$. We assume that paternal and maternal IGEs are equal, but we relate the results for this model to a model that allows for different paternal and maternal IGEs later.

In a random-mating population, the phenotypic variance can be decomposed as in the RDR method for estimating heritability [14] :

$$\text{Var}\left(Y_{ij}\right) = \text{Var}\left(\Delta_{ij}\right) + \text{Var}\left(\eta_{p(i)} + \eta_{m(i)}\right) + 2\,\text{Cov}\left(\Delta_{ij}, \eta_{p(i)} + \eta_{m(i)}\right) + \sigma_\epsilon^2;$$

where

$$v_g = \text{Var}\left(\Delta_{ij}\right) = 2\sum_{l=1}^{L} \delta_l^2 f_l \left(1 - f_l\right)$$

$$v_{e\sim g} = \text{Var}\left(\eta_{p(i)} + \eta_{m(i)}\right) = 4\sum_{l=1}^{L} \eta_l^2 f_l \left(1 - f_l\right);$$

$$c_{g,e} = 2\,\text{Cov}\left(\Delta_{ij}, \eta_{p(i)} + \eta_{m(i)}\right) = 4\sum_{l=1}^{L} \eta_l \delta_l f_l \left(1 - f_l\right)$$

so that

$$\mathrm{Var}(Y_{ij}) = v_g + v_{e\sim g} + c_{g,e} + \sigma_\epsilon^2.$$

Note that for this we have made the further assumption that the $L$ causal loci segregate independently so are uncorrelated in a random-mating population.

### 2 Assortative Mating Model

We consider assortative mating that has reached an equilibrium. From Chapter 4 of Crow and Kimura[1], the correlations between alleles at equilibrium are given in Figure S1.

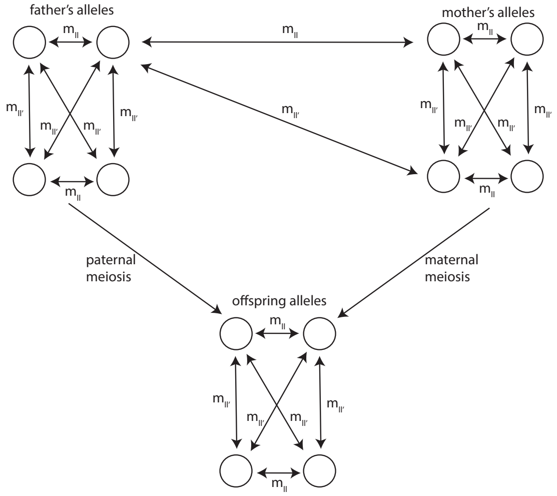


Fig. S1: Correlations between alleles in parents and offspring. Here, double-headed arrows indicate correlations between alleles. Assortative mating induces correlations between alleles in the mother and alleles in the father. At equilibrium, the correlations between alleles are as shown for two loci $l$ and $l'$[1]

We aim to express the equilibrium phenotypic variance in terms of the random-mating variance components (Equation 1) and the equilibrium correlations between the parents' direct and indirect components (Figure 1).

### 3 Equilibrium variance decomposition

We generalise the approach taken in Section 4.8 of Crow and Kimura[1] to apply to a trait determined by both DGEs and IGEs from parents.

#### 3.1 Equilibrium variance due to direct genetic effects

We have that:

$$\mathrm{Var}(\Delta_{ij}) = 2\sum_{l=1}^{L} \delta_l^2 f_l(1-f_l)[1+m_{ll}] + 4\sum_{l\neq l'} \delta_l \delta_{l'} \sqrt{f_l(1-f_l)f_{l'}(1-f_{l'})}m_{ll'}. \tag{1}$$

We relate this to $\mathrm{Cov}(\Delta_{m(i)}, \Delta_{p(i)}) = r_\delta \mathrm{Var}(\Delta_{ij})$, where $r_\delta = \mathrm{Corr}(\Delta_{m(i)}, \Delta_{p(i)})$ (Figure 1). We have that

$$\mathrm{Cov}(\Delta_{m(i)}, \Delta_{p(i)}) = 4\sum_{l,l'} \delta_l \delta_{l'} \sqrt{f_l(1-f_l)f_{l'}(1-f_{l'})}m_{ll'}. \tag{2}$$

We define the effective number of independently segregating loci of equal contribution to the variance due to DGEs, $L_\delta$:

$$L_\delta = \frac{\sum_{l,l'} \delta_l \delta_{l'} \sqrt{f_l(1-f_l)f_{l'}(1-f_{l'})}m_{ll'}}{\sum_{l=1}^{L} f_l(1-f_l)\delta_l^2 m_{ll}}. \tag{3}$$

Note that if all loci had equal DGEs and equal allele frequencies, then at equilibrium $m_{ll'} = m_{ll} = r \; \forall \; l, l'$, and therefore $L_\delta = L$. See Crow and Kimura section 4.7 for further details on the model with $L$ independently segregating loci with equal frequency and equal effects[1]. If many common genome-wide variants contribute to the variance explained by direct genetic effects, $L_\delta$ will be large. This allows us to express $\mathrm{Var}(\Delta_{ij})$ as

$$\mathrm{Var}(\Delta_{ij}) = v_g + r_\delta \mathrm{Var}(\Delta_{ij}) - r_\delta \frac{\mathrm{Var}(\Delta_{ij})}{2L_\delta}. \tag{4}$$

Let the equilibrium variance $\mathrm{Var}(\Delta_{ij})$ be $v_g^{\mathrm{eq}}$, then

$$v_g^{\mathrm{eq}} = \frac{v_g}{1-(1-\frac{1}{2L_\delta})r_\delta} \approx \frac{v_g}{1-r_\delta} \text{ for large } L_\delta. \tag{5}$$

#### 3.2 Equilibrium variance due to parental indirect genetic effects

As above in Subsection 3.1, it can be shown that at equilibrium

$$\mathrm{Var}(\eta_{p(i)}) = \mathrm{Var}(\eta_{m(i)}) = \frac{v_{e\sim g}}{2[1-(1-\frac{1}{2L_\eta})r_\eta]}, \tag{6}$$

where $r_\eta = \mathrm{Corr}(\eta_{p(i)}, \eta_{m(i)})$ and

$$L_\eta = \frac{\sum_{l,l'} \eta_l \eta_{l'} \sqrt{f_l(1-f_l)f_{l'}(1-f_{l'})}m_{ll'}}{\sum_{l=1}^{L} f_l(1-f_l)\eta_l^2 m_{ll}} \tag{7}$$

is the effective number of independently segregating loci of equal contribution to the variance due to parental IGEs.

Let $v^{\text{eq}}_{e\sim g}$ be the equilibrium variance due to parental IGEs. We have that

$$v^{\text{eq}}_{e\sim g} = \frac{v_{e\sim g}}{1-(1-\frac{1}{2L_\eta})r_\eta} + 2\text{Cov}(\eta_{p(i)}, \eta_{m(i)}). \tag{8}$$

We further have that

$$\text{Cov}(\eta_{p(i)}, \eta_{m(i)}) = r_\eta \frac{v_{e\sim g}}{2[1-(1-\frac{1}{2L_\eta})r_\eta]}. \tag{9}$$

Therefore,

$$v^{\text{eq}}_{e\sim g} = \frac{1+r_\eta}{1-(1-\frac{1}{2L_\eta})r_\eta} v_{e\sim g}. \tag{10}$$

#### 3.3 Equilibrium variance due to covariance between direct and indirect genetic effects

We compute the phenotypic variance due covariance between direct and indirect genetic effects:

$$c^{\text{eq}}_{g,e} = 2\text{Cov}(\Delta_{ij}, \eta_{p(i)} + \eta_{m(i)}); \tag{11}$$

$$= 4\text{Cov}(\Delta_{ij}, \eta_{p(i)}). \tag{12}$$

To compute $\text{Cov}(\Delta_{ij}, \eta_{p(i)})$, we use the fact that $\Delta_{ij} \perp \eta_{p(i)} | G_{\text{par}(i)}$, and $\mathbb{E}[\Delta_{ij}|G_{\text{par}(i)}] = (\Delta_{p(i)} + \Delta_{m(i)})/2$. Therefore,

$$\text{Cov}(\Delta_{ij}, \eta_{p(i)}) = \frac{1}{2}\left(\text{Cov}(\Delta_{p(i)}, \eta_{p(i)}) + \text{Cov}(\Delta_{p(i)}, \eta_{m(i)})\right); \tag{13}$$

$$= \frac{r^c_{\delta\eta} + r^\tau_{\delta\eta}}{2}\sqrt{\frac{v_g v_{e\sim g}}{2[1-(1-1/(2L_\delta))r_\delta][1-(1-1/(2L_\eta))r_\eta]}}. \tag{14}$$

Therefore,

$$c^{\text{eq}}_{g,e} = (r^c_{\delta\eta} + r^\tau_{\delta\eta})\sqrt{\frac{2v_g v_{e\sim g}}{[1-(1-1/(2L_\delta))r_\delta][1-(1-1/(2L_\eta))r_\eta]}}. \tag{15}$$

Letting $L_\delta \to \infty$ and $L_\eta \to \infty$, we obtain

$$c^{\text{eq}}_{g,e} = (r^c_{\delta\eta} + r^\tau_{\delta\eta})\sqrt{\frac{2v_g v_{e\sim g}}{(1-r_\delta)(1-r_\eta)}}. \tag{16}$$

##### 3.3.1 Relationship to random-mating variance component

We now relate the $c_{g,e}^{\text{eq}}$ to $c_{g,e}$ assuming that $c_{g,e} \neq 0$.

$$\text{Cov}(\Delta_{ij}, \eta_{p(i)}) = \sum_{l=1}^{L} \delta_l \eta_l f_l(1-f_l)[1+3m_{ll}] + 4\sum_{l \neq l'} \delta_l \eta_{l'} \sqrt{f_l(1-f_l)f_{l'}(1-f_{l'})} m_{ll'}. \tag{17}$$

This can be related to

$$\text{Cov}(\Delta_{m(i)}, \eta_{p(i)}) = 4\sum_{l,l'} \delta_l \eta_{l'} \sqrt{f_l(1-f_l)f_{l'}(1-f_{l'})} m_{ll'}. \tag{18}$$

We define

$$L_{\delta\eta} = \frac{\sum_{l,l'} \delta_l \eta_{l'} \sqrt{f_l(1-f_l)f_{l'}(1-f_{l'})} m_{ll'}}{\sum_{l=1}^{L} f_l(1-f_l)\delta_l \eta_l m_{ll}}, \tag{19}$$

the effective number of independent loci of equal contribution to the variance due to covariance between DGEs and IGEs. Note that if all loci had equal frequency and equal DGE and IGE, then $L_{\delta\eta} = L$. Assuming that $c_{g,e} \neq 0$, we have that

$$\text{Cov}(\Delta_{ij}, \eta_{p(i)}) = \frac{c_{g,e}}{4} + \left(1 - \frac{1}{4L_{\delta\eta}}\right) \text{Cov}(\Delta_{m(i)}, \eta_{p(i)}). \tag{20}$$

From above, we have that

$$\text{Cov}(\Delta_{ij}, \eta_{p(i)}) = \frac{r_{\delta\eta}^c + r_{\delta\eta}^\tau}{2} \sqrt{\text{Var}(\Delta_{ij})\text{Var}(\eta_{p(i)})}; \tag{21}$$

and

$$\text{Cov}(\Delta_{m(i)}, \eta_{p(i)}) = r_{\delta\eta}^\tau \sqrt{\text{Var}(\Delta_{ij})\text{Var}(\eta_{p(i)})}. \tag{22}$$

Therefore,

$$\text{Cov}(\Delta_{m(i)}, \eta_{p(i)}) = \frac{2r_{\delta\eta}^\tau}{r_{\delta\eta}^c + r_{\delta\eta}^\tau} \text{Cov}(\Delta_{ij}, \eta_{p(i)}), \tag{23}$$

and therefore

$$\text{Cov}(\Delta_{ij}, \eta_{p(i)}) = \frac{c_{g,e}}{4} + \left(1 - \frac{1}{4L_{\delta\eta}}\right) \frac{2r_{\delta\eta}^\tau}{r_{\delta\eta}^c + r_{\delta\eta}^\tau} \text{Cov}(\Delta_{ij}, \eta_{p(i)}). \tag{24}$$

After some rearrangement, it can be shown that

$$\text{Cov}(\Delta_{ij}, \eta_{p(i)}) = \frac{r_{\delta\eta}^c + r_{\delta\eta}^\tau}{r_{\delta\eta}^c - \left(1 - \frac{1}{2L_{\delta\eta}}\right) r_{\delta\eta}^\tau} \frac{c_{g,e}}{4}; \tag{25}$$

and therefore

$$c_{g,e}^{\text{eq}} = \frac{r_{\delta\eta}^c + r_{\delta\eta}^\tau}{r_{\delta\eta}^c - \left(1 - \frac{1}{2L_{\delta\eta}}\right) r_{\delta\eta}^\tau} c_{g,e}. \tag{26}$$

##### 3.3.2 Expression in terms of direct and indirect genetic effect correlation

We now derive the relationship between $r^c_{\delta\eta}$ and $r^\tau_{\delta\eta}$ to give an alternative expression for $c^{\mathrm{eq}}_{g,e}$. Recall that (Figure 1) $r^c_{\delta\eta}$ is the correlation between direct and indirect effect components within an individual:

$$r^c_{\delta\eta} = \frac{\mathrm{Cov}\left(\Delta_{p(i)}, \eta_{p(i)}\right)}{\sqrt{\mathrm{Var}\left(\Delta_{p(i)}\right)\mathrm{Var}\left(\eta_{p(i)}\right)}}$$

We now define the correlation between DGE and IGE components in an individual under random mating to be $r^0_{\delta\eta}$, which can be expressed in terms of the random-mating variance components:

$$r^0_{\delta\eta} = \frac{c_{g,e}}{\sqrt{2v_g v_{e\sim g}}}.$$

If we further assume equal allele frequencies, i.e. $f_l = f \,\forall\, l$, then this is simply the genome-wide correlation of the direct and indirect genetic effects:

$$r^0_{\delta\eta} = \frac{\sum_{l=1}^L \delta_l \eta_l}{\sqrt{\left(\sum_{l=1}^L \delta_l^2\right)\left(\sum_{l=1}^L \eta_l^2\right)}}.$$

Without assuming equal allele frequencies, it is the genome-wide correlation of the direct and indirect genetic effects for genotypes standardized to have variance 1 .

Now we compute $r^c_{\delta\eta}$ at equilibrium. First, we compute

$$\mathrm{Cov}\left(\Delta_{p(i)}, \eta_{p(i)}\right) = \sum_{l=1}^{L} \delta_l \eta_l 2 f_l\left(1-f_l\right)\left[1+m_{ll}\right] + 4\sum_{l \neq l'} \delta_l \eta_{l'} \sqrt{f_l\left(1-f_l\right) f_{l'}\left(1-f_{l'}\right)} m_{ll'}.$$

Following a similar procedure to above, we obtain:

$$\mathrm{Cov}\left(\Delta_{p(i)}, \eta_{p(i)}\right) = \frac{c_{g,e}}{2} + \left(1 - \frac{1}{2L_{\delta\eta}}\right)\mathrm{Cov}\left(\Delta_{m(i)}, \eta_{p(i)}\right)$$

Now we obtain the correlation at equilibrium:

$$r^c_{\delta\eta} = \frac{c_{g,e}}{2\sqrt{\mathrm{Var}\left(\Delta_{p(i)}\right)\mathrm{Var}\left(\eta_{p(i)}\right)}} + \left(1 - \frac{1}{2L_{\delta\eta}}\right)\frac{\mathrm{Cov}\left(\Delta_{m(i)}, \eta_{p(i)}\right)}{\sqrt{\mathrm{Var}\left(\Delta_{p(i)}\right)\mathrm{Var}\left(\eta_{p(i)}\right)}}$$

We can then express this in terms of $r^\tau_{\delta\eta} = \mathrm{Corr}\left(\Delta_{m(i)}, \eta_{p(i)}\right)$, $r_\delta$, $r_\eta$, and the random-mating variance components. Letting $L_{\delta\eta} \to \infty$,

$$r^c_{\delta\eta} = \frac{c_{g,e}/2}{\sqrt{\frac{v_g v_{e\sim g}}{2(1-r_\delta)(1-r_\eta)}}} + r^\tau_{\delta\eta}$$

This can then be further simplified by expression in terms of $r^0_{\delta\eta}$ :

$$r^c_{\delta\eta} = r^0_{\delta\eta}\sqrt{(1-r_\delta)(1-r_\eta)} + r^\tau_{\delta\eta}$$

We can then substitute this into the expression for $c^{\text{eq}}_{g,e}$ derived above:

$$c^{\text{eq}}_{g,e} = \left(1 + \frac{2}{\sqrt{(1-r_\delta)(1-r_\eta)}}\frac{r^\tau_{\delta\eta}}{r^0_{\delta\eta}}\right)c_{g,e}.$$

#### 3.4 Different maternal and paternal indirect genetic effects

If we allow the maternal and paternal indirect effects to be different, i.e.

$$\eta_{p(i)} = \sum_{l=1}^{L}\eta_{pl}\left(g_{p(i)l} - 2f_l\right); \eta_{m(i)} = \sum_{l=1}^{L}\eta_{ml}\left(g_{m(i)l} - 2f_l\right),$$

then we can write the phenotype model as (see[2])

$$Y_{ij} = \Delta_{ij} + \eta_{p(i)} + \eta_{m(i)} + \lambda_{p(i)} - \lambda_{m(i)} + \epsilon_{ij}$$

which is the same as in 1, except

$$\eta_l = (\eta_{pl} + \eta_{ml})/2; \lambda_{p(i)} = \sum_{l=1}^{L}\lambda_l g_{p(i)l}; \lambda_{m(i)} = \sum_{l=1}^{L}\lambda_l g_{m(i)l}; \lambda_l = (\eta_{pl} - \eta_{ml})/2.$$

Ignoring the term involving the differences between paternal and maternal indirect effects, this model is equivalent to the model considered above with $\eta_l$ replaced by $(\eta_{pl} + \eta_{ml})/2$, i.e. the average of paternal and maternal indirect effects.

Now we consider the statistical relationship between the difference term and the other non-residual terms. We first consider the covariance between the DGE component and the difference term. As the offspring genotype is equally related to maternal and paternal genotypes, i.e. by symmetry,

$$\text{Cov}\left(\Delta_{ij}, \lambda_{p(i)} - \lambda_{m(i)}\right) = 0.$$

We now consider the covariance between the sum and difference terms for parental IGEs. Since $\text{Var}\left(g_{p(i)l}\right) = \text{Var}\left(g_{m(i)l}\right)$,

$$\text{Cov}\left(g_{p(i)l} + g_{m(i)l}, g_{p(i)l} - g_{m(i)l}\right) = 0.$$

For distinct loci, we have that

$$\text{Cov}\left(g_{p(i)l} + g_{m(i)l}, g_{p(i)l'} - g_{m(i)l'}\right) = 2\left(4m_{ll'} - 4m_{ll'}\right) = 0.$$

Therefore,

$$\text{Cov}\left(\eta_{p(i)}+\eta_{m(i)},\lambda_{p(i)}-\lambda_{m(i)}\right)=0.$$

This shows that the difference component is uncorrelated with the DGE and average parental IGE components. Therefore, for many applications, the difference term can be subsumed into the residual and the variance decomposition for the model with equal paternal and maternal indirect effects used — along with the interpretation that the parental IGE is the average parental IGE.

However, for some applications, such as calculating covariances between relatives, the difference term remains important. We therefore give a variance decomposition at equilibrium including the difference term. First, we give the random-mating variance of $\lambda_{p(i)}-\lambda_{m(i)}$ :

$$v_{\lambda}=\text{Var}\left(\lambda_{p(i)}-\lambda_{m(i)}\right)=\sum_{l=1}^{L}\lambda_l^2 4f_l\left(1-f_l\right).$$

Now we consider that at equilibrium, $\text{Corr}\left(\lambda_{p(i)},\lambda_{m(i)}\right)=r_{\lambda}$. Let $v_{lp}^{\text{eq}}$ be the equilibrium variance of $\lambda_{p(i)}$, then the at equilibrium we have

$$\text{Var}\left(\lambda_{p(i)}-\lambda_{m(i)}\right)=2\left(1-r_{\lambda}\right)v_{lp}^{\text{eq}}.$$

Following a similar procedure to above for the other variance components, it can be shown that

$$v_{lp}^{\text{eq}}=\frac{v_{\lambda}}{2\left(1-\left(1-2/L_{\lambda}\right)r_{\lambda}\right)},$$

where

$$L_{\lambda}=\frac{\sum_{l,l'}\lambda_l\lambda_{l'}\sqrt{f_l\left(1-f_l\right)f_{l'}\left(1-f_{l'}\right)}4m_{ll'}}{\sum_{l=1}^{L}\lambda_l^2 f_l\left(1-f_l\right)m_{ll}}$$

Therefore, at equilibrium

$$\text{Var}\left(\lambda_{p(i)}-\lambda_{m(i)}\right)=\frac{1-r_{\lambda}}{1-\left(1-2/L_{\lambda}\right)r_{\lambda}}v_{\lambda}.$$

In limit as $L_{\lambda}\to\infty$,

$$\text{Var}\left(\lambda_{p(i)}-\lambda_{m(i)}\right)=v_{\lambda}.$$

This shows that assortative mating makes approximately no difference in the variance due to differences between paternal and maternal IGEs when there are many variants genome-wide contributing to differences between maternal and paternal IGE components. We can thereby generalize the equilibrium variance decomposition to include the variance component due to differences between maternal and paternal IGEs:

$$\mathrm{Var}\left(Y_{ij}\right)=\frac{v_g}{1-r_\delta}+\frac{1+r_\eta}{1-r_\eta}v_{e\sim g}+(r^c_{\delta\eta}+r^\tau_{\delta\eta})\sqrt{\frac{2v_gv_{e\sim g}}{(1-r_\delta)(1-r_\eta)}}+v_\lambda+\sigma^2_\epsilon.$$

#### 3.5 Primary phenotypic assortment

Previously, we did not consider the mechanism of assortment: just that we had reached an equilibrium where the correlations between alleles are constant across generations. Here we consider what the equilibrium correlation between parents' direct effect components would be under a model of assortative mating due to matching on the phenotype, called primary phenotypic assortment. The key assumption is that the paternal DGE and IGE components are conditionally independent of the maternal DGE and IGE components given the maternal and paternal phenotypes:

$$\Delta_{p(i)},\eta_{p(i)}\perp\Delta_{m(i)},\eta_{m(i)}\mid Y_{m(i)},Y_{p(i)}$$

Under this assumption and a further assumption that $\mathbb{E}\left[\Delta_{p(i)}\mid Y_{p(i)}\right]$ is a linear function of $Y_{p(i)}$ (see Nagylaki 1982[3] for a discussion of the conditions under which this assumption holds), we have that

$$\begin{aligned}r_\delta&=\frac{\mathrm{Cov}\left(\Delta_{p(i)},\Delta_{m(i)}\right)}{v^{\mathrm{eq}}_g};\\&=\frac{\mathrm{Cov}\left(\mathbb{E}\left[\Delta_{p(i)}\mid Y_{p(i)}\right],\mathbb{E}\left[\Delta_{m(i)}\mid Y_{m(i)}\right]\right)}{v^{\mathrm{eq}}_g};\\&=\frac{\mathrm{Cov}\left(\Delta_{p(i)},Y_{p(i)}\right)^2}{v^{\mathrm{eq}}_yv^{\mathrm{eq}}_g}r_Y;\end{aligned}$$

where $v^{\mathrm{eq}}_y$ is the equilibrium phenotypic variance. It is trivial to derive that $\mathrm{Cov}\left(\Delta_{p(i)},Y_{p(i)}\right)=v^{\mathrm{eq}}_g+c^{\mathrm{eq}}_{g,e}/2$, and therefore, after some rearrangement,

$$r_\delta=h^2_{\mathrm{eq}}r_Y\left[1+\frac{c^{\mathrm{eq}}_{g,e}}{v^{\mathrm{eq}}_g}\left(1+\frac{c^{\mathrm{eq}}_{g,e}}{4v^{\mathrm{eq}}_g}\right)\right],$$

We therefore see that parental IGEs, when correlated with DGEs, can inflate the equilibrium correlation between DGE components of parents over what would be expected from DGEs alone. The same logic implies that the other correlations in Figure 1, and therefore the equilibrium phenotypic variance will be further inflated by AM when DGEs and IGEs are correlated.

### 4 Estimating heritability using realized relatedness

Here, we examine heritability estimation using realized relatedness between siblings ('sib-regression'), as first proposed in 2006 by Visscher et al.[4]. Let $R_{ijk}$ be the realized relatedness between sibling $j$ and sibling $k$ in family $i$. While the expected relatedness between siblings is given by the pedigree, the realized relatedness varies around the expectation due to random segregations during meiosis in the parents, leading to variation in the fractions of the genome the siblings share identical-by-descent (IBD). We first derive the phenotypic covariance between siblings in terms of their realized relatedness, then we derive the bias due to AM in sib-regression estimates of heritability.

#### 4.1 Equilibrium covariance between siblings

We show how to express the equilibrium phenotypic covariance between siblings in terms of their realized relatedness and the equilibrium decomposition of the phenotypic variance. The covariance between the DGE components of a sibling pair is:

$$\text{Cov}\left(\Delta_{i1}, \Delta_{i2}\right) = \sum_{l=1}^{L} \delta_l^2 \,\text{Cov}\left(g_{i1l}, g_{i2l}\right) + 4 \sum_{l \neq l'} \delta_l \delta_{l'} \sqrt{f_l\left(1-f_l\right) f_{l'}\left(1-f_{l'}\right)} m_{ll'}$$

The covariance between siblings' genotypes conditional on the IBD state at the locus is:

$$\text{Cov}\left(g_{i1l}, g_{i2l}\right) = \begin{cases} 4 m_{ll} f_l\left(1-f_l\right), \text{IBD}_0 \\ \left(1+3m_{ll}\right) f_l\left(1-f_l\right), \text{IBD}_1 \\ 2\left(1+m_{ll}\right) f_l\left(1-f_l\right), \text{IBD}_2 \end{cases}$$

Let $P_0, P_1, P_2$ be the proportion of the genome shared in IBD states 0, 1, and 2, respectively. Therefore, assuming causal variants are located at random with respect to IBD sharing,

$$\text{Cov}\left(g_{i1l}, g_{i2l}\right) = f_l\left(1-f_l\right)\left[P_1 + 2P_2 + m_{ll}\left(4P_0 + 3P_1 + 2P_2\right)\right].$$

The realised relatedness between the siblings is $R_{ijk} = \left(P_1 + 2P_2\right)/2$. Therefore,

$$\text{Cov}\left(g_{i1l}, g_{i2l}\right) = 2f_l\left(1-f_l\right)\left[R_{ijk} + m_{ll}\left(2P_0 + (3/2)P_1 + P_2\right)\right].$$

Since $P_0 + P_1 + P_2 = 1, 2P_0 + (3/2)P_1 + P_2 = 2 - R_{ijk}$. Therefore,

$$\text{Cov}\left(g_{i1l}, g_{i2l}\right) = 2f_l\left(1-f_l\right)\left[R_{ijk} + (2 - R_{ijk}) m_{ll}\right].$$

This gives the covariance between siblings' DGE components as

$$\text{Cov}\left(\Delta_{i1}, \Delta_{i2}\right) = v_g R_{ijk} + 2(2 - R_{ijk}) \sum_{l=1}^{L} \delta_l^2 f_l\left(1-f_l\right) m_{ll} + 4 \sum_{l \neq l'} \delta_l \delta_{l'} \sqrt{f_l\left(1-f_l\right) f_{l'}\left(1-f_{l'}\right)} m_{ll'}.$$

We now relate this to $\text{Cov}\left(\Delta_{m(i)}, \Delta_{p(i)}\right) = r_\delta v_g^{\text{eq}}$ :

$$\text{Cov}\left(\Delta_{i1}, \Delta_{i2}\right) = v_g R_{ijk} + 2(2 - R_{ijk}) \sum_{l=1}^{L} \delta_l^2 f_l\left(1-f_l\right) m_{ll} + r_\delta v_g^{\text{eq}} - \sum_{l=1}^{L} \delta_l^2 f_l\left(1-f_l\right) m_{ll}.$$

Now expressing in terms of $L_\delta$ :

$$\text{Cov}\left(\Delta_{i1}, \Delta_{i2}\right) = v_g R_{ijk} + \left(1 + \frac{3 - 2R_{ijk}}{4L_\delta}\right) r_\delta v_g^{\text{eq}}.$$

Now letting $L_\delta \to \infty$,

$$\text{Cov}\left(\Delta_{i1}, \Delta_{i2}\right) \to v_g R_{ijk} + r_\delta v_g^{\text{eq}}.$$

We therefore see that the realized relatedness does not track the assortative mating induced inflation of the variance due to direct effects. Setting $R_{ijk} = 0.5$, the expected relatedness given the pedigree, we obtain

$$\text{Cov}\left(\Delta_{i1}, \Delta_{i2}\right) = \frac{1 + r_\delta}{2} v_g^{\text{eq}},$$

the same as based on pedigree alone. This gives the phenotypic covariance between siblings as

$$\text{Cov}\left(Y_{i1}, Y_{i2}\right) = v_g R_{ijk} + r_\delta v_g^{\text{eq}} + v_{e\sim g}^{\text{eq}} + c_{g,e}^{\text{eq}} + \text{Cov}\left(\epsilon_{i1}, \epsilon_{i2}\right).$$

Allowing for finite $L_\delta$, we obtain

$$\text{Cov}\left(Y_{ij}, Y_{ik}\right) = \left(1 - \frac{r_\delta}{2L_\delta - \left(2L_\delta - 1\right) r_\delta}\right) v_g R_{ijk} + r_\delta v_g^{\text{eq}} + v_{e\sim g}^{\text{eq}} + c_{g,e}^{\text{eq}} + \text{Cov}\left(\epsilon_{ij}, \epsilon_{ik}\right)$$

#### 4.2 Estimating heritability by sib-regression

Heritability is estimated by the slope of the regression of $\left(Y_{ij} - \mu_y\right)\left(Y_{ik} - \mu_y\right)/v_y^{\text{eq}}$ onto $R_{ijk}$ across sibling pairs[4, 5], where $\mu_y$ is the phenotypic mean and $v_y^{\text{eq}}$ is the equilibrium phenotypic variance. Since

$$\mathbb{E}\left[\left(Y_{ij} - \mu_y\right)\left(Y_{ik} - \mu_y\right)/v_y^{\text{eq}} \mid R_{ijk}\right] = \left(r_\delta v_g^{\text{eq}} + v_{e\sim g}^{\text{eq}} + c_{g,e}^{\text{eq}} + \mathbb{E}[\epsilon_{ij}\epsilon_{ik}]\right)/v_y^{\text{eq}} + \left(1 - \frac{r_\delta}{2L_\delta - \left(2L_\delta - 1\right) r_\delta}\right) \frac{v_g}{v_y^{\text{eq}}} R_{ijk}.$$

Assuming that $\epsilon_{ij}\epsilon_{ik}$ is uncorrelated with $R_{ijk}$ across sibling pairs (which is violated when there are indirect genetic effects between siblings[5]), this implies that performing the regression across siblings gives as slope

$$\left(1 - \frac{r_\delta}{2L_\delta - \left(2L_\delta - 1\right) r_\delta}\right) h_f^2 \to h_f^2 = \frac{v_g}{v_y^{\text{eq}}} \text{ as } L_\delta \to \infty$$

i.e. the random mating variance of the DGE component divided by the equilibrium phenotypic variance. This is smaller than the equilibrium heritability, $h^2_{\text{eq}}$, by a factor of $v_g/v_g^{\text{eq}} = (1 - r_\delta)$. Although unlikely to be relevant for complex traits in humans, the above result implies that there would be a further downward bias when there is AM for a phenotype affected by only a small (effective) number of loci. For example, if $L_\delta = 1$, as for a phenotype affected by a single variant, then the heritability estimate would be

$$\left(1 - \frac{r_\delta}{2 - r_\delta}\right) h_f^2,$$

which approaches zero as $r_\delta$ approaches 1.

The intercept is

$$r_\delta h_{\text{eq}}^2 + \left(v_{e\sim g}^{\text{eq}} + c_{g,e}^{\text{eq}} + \mathbb{E}[\epsilon_{ij}\epsilon_{ik}]\right)/v_y^{\text{eq}}, \tag{27}$$

where the $r_\delta h_{\text{eq}}^2$ term captures the increased correlation between siblings' DGE components due to AM, and the remaining terms capture variance explained by environmental factors shared between siblings.

This result (in the limit as $L_\delta \to \infty$) agrees with the hypothesis put forward in Kemper et al.[10], which argued that sib-regression estimates the random mating genetic variance divided by the phenotypic variance in the present generation, which is $h_f^2$ at equilibrium. Kemper et al. supported their argument through a theoretical derivation and simulations of one generation of assortative mating. We show here that their theoretical argument was incorrect. In Supplementary Note Section 3.2 of Kemper et al. (pages 60-61 of the supplement) they use the following result to argue that sib-regression estimates $h_f^2$:

$$\mathbb{E}[\delta_{i1}\delta_{i2}|R_{ijk}] = v_g R_{ijk}, \tag{28}$$

which is stated without proof. However, as we prove above (Equation 4.1),

$$\mathbb{E}[\delta_{i1}\delta_{i2}|R_{ijk}] = v_g R_{ijk} + \left(1 + \frac{3 - 2R_{ijk}}{4L_\delta}\right) r_\delta v_g^{\text{eq}} \to v_g R_{ijk} + r_\delta v_g^{\text{eq}} \text{ as } L_\delta \to \infty. \tag{29}$$

However, using the incorrect result $\mathbb{E}[\delta_{i1}\delta_{i2}|R_{ijk}] = v_g R_{ijk}$ in their derivation still gives the correct slope for sib-regression because the the missing $r_\delta v_g^{\text{eq}}$ term is uncorrelated with $R_{ijk}$ across sibling pairs. The result they give for the slope in the section is also missing a $r_\delta v_g^{\text{eq}}$ term since they ignore the inflation of genetic covariance between siblings due to AM when giving the phenotypic covariance between siblings.

### 5 Polygenic index analysis under random-mating

Here we derive the expected regression coefficients from two-generation PGI analysis under random-mating in terms of the weight vector of the PGI. Let

$$\text{PGI}_{ij} = \frac{1}{\sqrt{v}} \sum_{l=1}^{L} w_l (g_{ijl} - 2f_l);\ \text{PGI}_{\text{par}(i)} = \frac{1}{\sqrt{v}} \sum_{l=1}^{L} w_l (g_{p(i)} + g_{m(i)} - 4f_l);. \tag{30}$$

where

$$v = \sum_{l=1}^{L} w_l^2 2f_l(1-f_l). \tag{31}$$

We consider performing the regression defined by:

$$Y_{ij} = \delta_{\text{PGI}} \text{PGI}_{ij} + \alpha_{\text{PGI}} \text{PGI}_{\text{par}(i)} + \epsilon_{ij}. \tag{32}$$

Let

$$\mathbf{X}_{ij} = \begin{bmatrix} \text{PGI}_{ij} \\ \text{PGI}_{\text{par}(i)} \end{bmatrix}.$$

Then we have that, under random-mating,

$$\text{Var}\left(\mathbf{X}_{ij}\right) = \begin{bmatrix} 1 & 1 \\ 1 & 2 \end{bmatrix};$$

and

$$\text{Var}\left(\mathbf{X}_{ij}\right)^{-1} = \begin{bmatrix} 2 & -1 \\ -1 & 1 \end{bmatrix}.$$

We also have that

$$\text{Cov}\left(\mathbf{X}_{ij}, Y_{ij}\right) = \frac{1}{\sqrt{v}} \begin{bmatrix} \sum_{l=1}^{L} \omega_l \delta_l 2f_l\left(1-f_l\right) + \sum_{l=1}^{L} \omega_l \eta_l 2f_l\left(1-f_l\right) \\ \sum_{l=1}^{L} \omega_l \delta_l 2f_l\left(1-f_l\right) + \sum_{l=1}^{L} \omega_l \eta_l 4f_l\left(1-f_l\right) \end{bmatrix}.$$

Therefore,

$$\begin{bmatrix} \delta_{\text{PGI}} \\ \alpha_{\text{PGI}} \end{bmatrix} = \text{Var}\left(\mathbf{X}_{ij}\right)^{-1} \text{Cov}\left(\mathbf{X}_{ij}, Y_{ij}\right) = \frac{1}{\sqrt{v}} \begin{bmatrix} \sum_{l=1}^{L} \omega_l \delta_l 2f_l\left(1-f_l\right) \\ \sum_{l=1}^{L} \omega_l \eta_l 2f_l\left(1-f_l\right) \end{bmatrix}.$$

Under random-mating, we also have that

$$\beta_{\text{PGI}} = \frac{\text{Cov}(Y_{ij}, \text{PGI}_{ij})}{\text{Var}(\text{PGI}_{ij})} = \delta_{\text{PGI}} + \alpha_{\text{PGI}}. \tag{33}$$

### 6 Direct genetic effect PGI at equilibrium

Here we derive the expected regression coefficients from two-generation PGI analysis of the true DGE PGI under assortative mating at equilibrium. This is defined by the following regression:

$$Y_{ij} = \delta_\delta \Delta_{ij} + \alpha_\delta \Delta_{\text{par}(i)} + \epsilon_{ij}; \tag{34}$$

where (Equation 1 and Figure 1)

$$\Delta_{ij} = \sum_{l=1}^{L} \delta_l (g_{ijl} - 2f_l);\ \ \Delta_{\text{par}(i)} = \sum_{l=1}^{L} \delta_l (g_{p(i)} + g_{m(i)} - 4f_l) = \Delta_{p(i)} + \Delta_{m(i)}. \tag{35}$$

Let

$$\mathbf{X}_{ij} = \begin{bmatrix} \Delta_{ij} \\ \Delta_{\text{par}(i)} \end{bmatrix}.$$

Then we have that

$$\text{Var}\left(\mathbf{X}_{ij}\right) = v_g^{\text{eq}} \begin{bmatrix} 1 & 1 + r_\delta \\ 1 + r_\delta & 2\left(1 + r_\delta\right) \end{bmatrix};$$

and

$$\text{Var}\left(\mathbf{X}_{ij}\right)^{-1} = \frac{1}{v_g^{\text{eq}}\left(1 - r_\delta^2\right)} \begin{bmatrix} 2\left(1 + r_\delta\right) & -\left(1 + r_\delta\right) \\ -\left(1 + r_\delta\right) & 1 \end{bmatrix}. \tag{36}$$

We also have that

$$\text{Cov}\left(\mathbf{X}_{ij}, Y_{ij}\right) = \begin{bmatrix} v_g^{\text{eq}} + c_{g,e}^{\text{eq}}/2 \\ \left(1 + r_\delta\right) v_g^{\text{eq}} + \text{Cov}\left(\Delta_{\text{par}(i)}, \eta_{p(i)} + \eta_{m(i)}\right) \end{bmatrix},$$

where we have used the fact that $\text{Cov}\left(\Delta_{ij}, \eta_{p(i)} + \eta_{m(i)}\right) = c_{g,e}^{\text{eq}}/2$. We can again use this to compute $\text{Cov}\left(\Delta_{\text{par}(i)}, \eta_{p(i)} + \eta_{m(i)}\right)$. Because $\Delta_{ij} \perp \eta_{p(i)}, \eta_{m(i)} \mid G_{\text{par}(i)}$ and $\mathbb{E}\left[\Delta_{ij} \mid G_{\text{par}(i)}\right] = \Delta_{\text{par}(i)}/2$,

$$\text{Cov}\left(\Delta_{ij}, \eta_{p(i)} + \eta_{m(i)}\right) = \text{Cov}\left(\Delta_{\text{par}(i)}/2, \eta_{p(i)} + \eta_{m(i)}\right);$$
$$\Rightarrow \text{Cov}\left(\Delta_{\text{par}(i)}, \eta_{p(i)} + \eta_{m(i)}\right) = c_{g,e}^{\text{eq}}.$$

Therefore,

$$\text{Cov}\left(\mathbf{X}_{ij}, Y_{ij}\right) = \begin{bmatrix} v_g^{\text{eq}} + c_{g,e}^{\text{eq}}/2 \\ \left(1 + r_\delta\right) v_g^{\text{eq}} + c_{g,e}^{\text{eq}} \end{bmatrix};$$

and therefore

$$\begin{bmatrix} \delta_\delta \\ \alpha_\delta \end{bmatrix} = \text{Var}\left(\mathbf{X}_{ij}\right)^{-1} \text{Cov}\left(\mathbf{X}_{ij}, Y_{ij}\right) = \begin{bmatrix} 1 \\ \frac{c_{g,e}^{\text{eq}}}{2(1+r_\delta)v_g^{\text{eq}}} \end{bmatrix}.$$

#### 6.1 Variance explained by parent and offspring DGE PGIs

We now compute the phenotypic variance explained the regression of parent and offspring DGE PGIs (Equation 34):

$$\text{Var}(Y_{ij}) = \text{Var}(\Delta_{ij}) + \alpha_\delta^2 \text{Var}(\Delta_{\text{par}(i)}) + 2\alpha_\delta \text{Cov}(\Delta_{ij}, \Delta_{\text{par}(i)}) + \sigma_\epsilon^2 \tag{37}$$
$$= v_g^{\text{eq}}[1 + 2\alpha_\delta^2(1 + r_\delta) + 2\alpha_\delta(1 + r_\delta)] + \sigma_\epsilon^2 \tag{38}$$
$$= v_g^{\text{eq}}[1 + 2(1 + r_\delta)\alpha_\delta(1 + \alpha_\delta)] + \sigma_\epsilon^2. \tag{39}$$

Therefore, the fraction of phenotypic variance explained by the regression is:

$$\frac{\text{Var}(\Delta_{ij} + \alpha_\delta \Delta_{\text{par}(i)})}{v_y^{\text{eq}}} = h_{\text{eq}}^2[1 + 2(1 + r_\delta)\alpha_\delta(1 + \alpha_\delta)]. \tag{40}$$

The increase in variance explained compared to if there were no IGEs (i.e. $\alpha_\delta = 0$) is

$$v_{\eta:\delta} \stackrel{\text{def}}{=} h_{\text{eq}}^2[1 + 2(1 + r_\delta)\alpha_\delta(1 + \alpha_\delta)] - h_{\text{eq}}^2 = 2(1 + r_\delta)\alpha_\delta(1 + \alpha_\delta)h_{\text{eq}}^2. \tag{41}$$

### 7 Incomplete direct genetic effect PGI

Here we derive results for two-generation analysis of an incomplete DGE PGI. We assume that all causal variants have equal allele frequency, $f$, and equal DGE, $\delta$. For some $0 < k \leq 1$, the incomplete direct DGE PGI for individual $j$ from family $i$ is:

$$\text{PGI}_{ij}^{\delta_k} = \delta \sum_{l=1}^{kL} (g_{ijl} - 2f), \tag{42}$$

and the equivalent incomplete parental DGE PGIs are:

$$\text{PGI}_{p(i)}^{\delta_k} = \delta \sum_{l=1}^{kL} (g_{p(i)l} - 2f);\ \text{PGI}_{m(i)}^{\delta_k} = \delta \sum_{l=1}^{kL} (g_{m(i)l} - 2f);\ \text{PGI}_{\text{par}(i)}^{\delta_k} = \text{PGI}_{p(i)}^{\delta_k} + \text{PGI}_{m(i)}^{\delta_k}. \tag{43}$$

#### 7.1 Equilibrium variance of incomplete direct genetic effect PGI

As above, we consider assortative mating to have reached an equilibrium with

$$\text{Corr}\left(\Delta_{p(i)}, \Delta_{m(i)}\right) = r_\delta$$

In this simplified model, the correlations between distinct alleles (Figure S1) are all equal, with

$$m_{ll} = m_{ll'} = \frac{r_\delta}{2L\,(1 - r_\delta) + r_\delta}$$

as first given by Sewall Wright in 1921[6]. (We also have that $L_\delta = L$ in this simplified model.)

First, we consider the equilibrium variance of $\text{PGI}_{ij}^{\delta_k}$ :

$$\begin{aligned}
\text{Var}\left(\text{PGI}_{ij}^{\delta_k}\right) &= \delta^2 kL2f(1-f)[1 + (2kL - 1)m] \\
&= kv_g[1 + (2kL - 1)m] \\
&= kv_g \frac{1 - (1-k)r_\delta}{1 - r_\delta + r_\delta/(2L)} \\
&\to kv_g \frac{1 - (1-k)r_\delta}{1 - r_\delta} \text{ as } L \to \infty.
\end{aligned}$$

The variance of the incomplete DGE PGI is inflated from $kv_g$ (under random-mating) to

$$\text{Var}\left(\text{PGI}_{ij}^{\delta_k}\right) \to kv_g \frac{1}{1 - r_k};$$

where

$$r_k = \text{Corr}\left(\text{PGI}_{p(i)}^{\delta_k}, \text{PGI}_{m(i)}^{\delta_k}\right).$$

#### 7.2 Relationship between correlations of incomplete and true DGE PGIs

By equating the two different expressions for the equilibrium variance of the incomplete PGI (above),

$$\mathrm{Var}\left(\mathrm{PGI}_{ij}^{\delta_k}\right) = kv_g \frac{1}{1-r_k} = kv_g \frac{1-(1-k)r_\delta}{1-r_\delta}, \tag{44}$$

we obtain the relationship between the equilibrium correlation between parents' DGE components, $r_\delta$, and the equilibrium correlation between parents' incomplete DGE PGIs, $r_k$:

$$r_k = \frac{kr_\delta}{1-(1-k)r_\delta}; r_\delta = \frac{r_k}{k+(1-k)r_k}.$$

#### 7.3 Direct to population effect ratio without indirect genetic effects

We assume a model without IGEs, i.e. $\eta_{p(i)} = \eta_{m(i)} = 0$, but with assortative mating at equilibrium. We first derive the 'population effect' of the incomplete DGE PGI, which is the equilibrium regression coefficient of $Y_{ij}$ on $\mathrm{PGI}_{ij}^{\delta_k}$.

The covariance between phenotype and incomplete DGE PGI is:

$$\mathrm{Cov}\left(\mathrm{PGI}_{ij}^{\delta_k}, Y_i\right) = \mathrm{Var}\left(\mathrm{PGI}_{ij}^{\delta_k}\right) + \delta^2 L^2 k(1-k)4mf(1-f).$$

Therefore, the population effect is:

$$\beta_{\mathrm{PGI}:k} = \frac{\mathrm{Cov}\left(\mathrm{PGI}_{ij}^{\delta_k}, Y_i\right)}{\mathrm{Var}\left(\mathrm{PGI}_{ij}^{\delta_k}\right)} = 1 + \frac{2(1-k)Lm}{1+(2kL-1)m};$$
$$= \frac{1+(2L-1)m}{1+(2kL-1)m};$$

where we have substituted in $\mathrm{Var}\left(\mathrm{PGI}_{ij}^{\delta_k}\right) = \delta^2 kL2f(1-f)[1+(2kL-1)m]$ from above. We now express $1+(2kL-1)m$ in terms of $r_\delta$ and $L$ :

$$1+(2kL-1)m = \frac{2L-(1-k)2Lr_\delta}{2L(1-r_\delta)+r_\delta}.$$

By setting $k=1$, we obtain:

$$1+(2L-1)m = \frac{2L}{2L(1-r_\delta)+r_\delta}$$

and therefore

$$\beta_{\text{PGI}:k} = \frac{\text{Cov}\left(\text{PGI}_{ij}^{\delta_k}, Y_i\right)}{\text{Var}\left(\text{PGI}_{ij}^{\delta_k}\right)} = \frac{2L}{2L - (1-k)2Lr_\delta};$$

$$= \frac{1}{1-(1-k)r_\delta} = 1 + \left(\frac{1}{k} - 1\right) r_k.$$

Since the direct effect of the incomplete DGE PGI is 1, we therefore have that

$$\frac{\delta_{\text{PGI}:k}}{\beta_{\text{PGI}:k}} = 1 - (1-k)r_\delta. \tag{45}$$

##### 7.3.1 Variance explained by incomplete DGE PGI

The variance explained by regression of phenotype onto incomplete DGE PGI is therefore:

$$\beta_{\text{PGI}:k}^2 \text{Var}\left(\text{PGI}_{ij}^{\delta_k}\right) = \left[1 + \left(\frac{1}{k} - 1\right) r_k \left(2 + r_k\right)\right] \frac{kv_g}{1 - r_k}.$$

We can compare this to the equilibrium genetic variance, $v_g^{\text{eq}} = v_g / (1 - r_\delta)$, to obtain the fraction of heritability explained by regression of phenotype onto incomplete PGI at equilibrium:

$$\frac{\beta_{\text{PGI}:k}^2 \text{Var}\left(\text{PGI}_{ij}^{\delta_k}\right)}{v_g^{\text{eq}}} = \left[1 + \left(\frac{1}{k} - 1\right) r_k \left(2 + r_k\right)\right] \frac{k}{1 + (1/k - 1)r_k}; \tag{46}$$

$$= \left[1 + \frac{(1/k-1)r_k\left(1+r_k\right)}{1+(1/k-1)r_k}\right] k; \tag{47}$$

$$= \left[1 + (1-k)r_\delta\left(1 + r_k\right)\right] k. \tag{48}$$

To obtain the fraction of phenotypic variance explained, we multiply by $h_{\text{eq}}^2$:

$$\frac{\beta_{\text{PGI}:k}^2 \text{Var}\left(\text{PGI}_{ij}^{\delta_k}\right)}{v_y^{\text{eq}}} = \left[1 + (1-k)r_\delta\left(1 + r_k\right)\right] kh_{\text{eq}}^2. \tag{49}$$

#### 7.4 Direct to population effect ratio with indirect effects

Consider the regression of phenotype onto offspring and parental true DGE PGIs outlined in the main text and in Section 6:

$$Y_{ij} = \Delta_{ij} + \alpha_\delta \Delta_{\text{par}(i)} + \epsilon_{ij}, \tag{50}$$

where

$$\alpha_\delta = \frac{c_{g,e}^{\text{eq}}}{2\left(1 + r_\delta\right) v_g^{\text{eq}}} \tag{51}$$

We now compute the population effect of the incomplete DGE PGI in terms of the parameters of the regression on the true DGE PGI (Equation 50):

$$\beta_{\text{PGI}:k} = \frac{\text{Cov}\left(\text{PGI}_{ij}^{\delta_k}, Y_i\right)}{\text{Var}\left(\text{PGI}_{ij}^{\delta_k}\right)} = \frac{1}{1-(1-k)r_\delta} + \alpha_\delta \frac{\text{Cov}\left(\text{PGI}_{ij}^{\delta_k}, \Delta_{\text{par}(i)}\right)}{\text{Var}\left(\text{PGI}_{ij}^{\delta_k}\right)}$$

where the first term is the population effect of the incomplete DGE PGI when there are no IGEs (above).

We now compute $\text{Cov}\left(\text{PGI}_{ij}^{\delta_k}, \Delta_{\text{par}(i)}\right)$. Since $\text{PGI}_{ij}^{\delta_k} \perp \Delta_{\text{par}(i)} | G_{\text{par}(i)}$ and $\mathbb{E}\left[\text{PGI}_{ij}^{\delta_k} \mid G_{\text{par}(i)}\right] = \left(\text{PGI}_{p(i)}^{\delta_k} + \text{PGI}_{m(i)}^{\delta_k}\right)/2$, we have

$$\begin{aligned}
\text{Cov}\left(\text{PGI}_{ij}^{\delta_k}, \Delta_{\text{par}(i)}\right) &= \text{Cov}\left(\mathbb{E}\left[\text{PGI}_{ij}^{\delta_k} \mid G_{\text{par}(i)}\right], \Delta_{\text{par}(i)}\right) \\
&= \frac{1}{2}\text{Cov}\left(\text{PGI}_{p(i)}^{\delta_k} + \text{PGI}_{m(i)}^{\delta_k}, \Delta_{p(i)} + \Delta_{m(i)}\right); \\
&= \text{Cov}\left(\text{PGI}_{p(i)}^{\delta_k}, \Delta_{p(i)}\right) + \text{Cov}\left(\text{PGI}_{p(i)}^{\delta_k}, \Delta_{m(i)}\right).
\end{aligned}$$

As we are at equilibrium, $\text{Cov}\left(\text{PGI}_{p(i)}^{\delta_k}, \Delta_{p(i)}\right) / \text{Var}\left(\text{PGI}_{ij}^{\delta_k}\right) = [1-(1-k)r_\delta]^{-1}$, as derived above for the offspring PGI. Therefore,

$$\frac{\text{Cov}\left(\text{PGI}_{ij}^{\delta_k}, \Delta_{\text{par}(i)}\right)}{\text{Var}\left(\text{PGI}_{ij}^{\delta_k}\right)} = \frac{1}{1-(1-k)r_\delta} + \frac{\text{Cov}\left(\text{PGI}_{p(i)}^{\delta_k}, \Delta_{m(i)}\right)}{\text{Var}\left(\text{PGI}_{ij}^{\delta_k}\right)}$$

Now we compute

$$\text{Cov}\left(\text{PGI}_{p(i)}^{\delta_k}, \Delta_{m(i)}\right) = \text{Cov}\left(\sum_{l=1}^{kL} \delta g_{p(i)l}, \sum_{l=1}^{L} \delta g_{m(i)l}\right) = \delta^2 k L^2 4mf(1-f).$$

Letting $k = 1$, we also have that

$$\text{Cov}\left(\Delta_{p(i)}, \Delta_{m(i)}\right) = \delta^2 L^2 4mf(1-f) = r_\delta v_g^{\text{eq}}$$

and therefore

$$\text{Cov}\left(\text{PGI}_{p(i)}^{\delta_k}, \Delta_{m(i)}\right) = k r_\delta v_g^{\text{eq}}$$

Therefore,

$$\frac{\text{Cov}\left(\text{PGI}_{p(i)}^{\delta_k}, \Delta_{m(i)}\right)}{\text{Var}\left(\text{PGI}_{ij}^{\delta_k}\right)} = \frac{k r_\delta v_g^{\text{eq}}}{k v_g^{\text{eq}}(1-(1-k)r_\delta)} = \frac{r_\delta}{1-(1-k)r_\delta}, \tag{52}$$

where we have substituted in $\text{Var}\left(\text{PGI}_{ij}^{\delta_k}\right) = kv_g^{\text{eq}}(1-(1-k)r_\delta)$ from Equation 44. This gives

$$\frac{\text{Cov}\left(\text{PGI}_{ij}^{\delta_k}, \Delta_{\text{par}(i)}\right)}{\text{Var}\left(\text{PGI}_{ij}^{\delta_k}\right)} = \frac{1+r_\delta}{1-(1-k)r_\delta}$$

Therefore,

$$\beta_{\text{PGI}:k} = \frac{1+(1+r_\delta)\alpha_\delta}{1-(1-k)r_\delta}.$$

Since the direct effect of the incomplete DGE PGI is 1, we obtain

$$\frac{\delta_{\text{PGI}:k}}{\beta_{\text{PGI}:k}} = \frac{1-(1-k)r_\delta}{1+(1+r_\delta)\alpha_\delta}. \tag{53}$$

##### 7.5 Average NTC of incomplete DGE PGI

We now compute the average NTC of the incomplete DGE PGI, $\alpha_{\text{PGI}:k}$. At equilibrium,

$$\beta_{\text{PGI}:k} = \delta_{\text{PGI}:k} + (1+r_k)\alpha_{\text{PGI}:k} = 1 + (1+r_k)\alpha_{\text{PGI}:k}. \tag{54}$$

Therefore,

$$\alpha_{\text{PGI}:k} = (\beta_{\text{PGI}:k} - 1)/(1+r_k) \tag{55}$$

$$= \frac{(1+r_\delta)\alpha_\delta + (1-k)r_\delta}{1+r_k}. \tag{56}$$

Since $\delta_{\text{PGI}:k} = 1$ and $\rho_k = 1-(1-k)r_\delta$, we also have

$$\frac{\alpha_{\text{PGI}:k}}{\delta_{\text{PGI}:k}} = \frac{(1+r_\delta)\alpha_\delta + (1-\rho_k)}{1+r_k}. \tag{57}$$

#### 8 Estimating $k$

For the inference procedure, one first needs to estimate $k$, the fraction of heritability the PGI would explain in a random-mating population. Consider that one has performed the following regression:

$$\frac{Y_{ij}}{\sqrt{v_y^{\text{eq}}}} = \delta_{\text{PGI}:k}\text{PGI}_{ij}^{\delta_k} + \alpha_{\text{PGI}:k}\text{PGI}_{\text{par}(i)}^{\delta_k} + \epsilon_{ij}, \tag{58}$$

where the proband and parental PGIs have been scaled by the inverse of the standard deviation of the proband PGI, so that the proband PGI has variance 1. (This differs from the theoretical derivations above that use the un-normalized incomplete DGE PGI.) The expected direct effect at equilibrium is (Equation 44):

$$\delta_{\text{PGI}:k} = \sqrt{\frac{kv_g}{v_y^{\text{eq}}(1-r_k)}}.$$

Therefore, the fraction of variance explained by the PGI in a random-mating population is

$$k = (1-r_k)\,\delta^2_{\text{PGI}:k}\frac{v_y^{\text{eq}}}{v_g} = \frac{(1-r_k)\delta^2_{\text{PGI}:k}}{h_f^2}.$$

To estimate $k$, consider that we have unbiased and statistically independent estimates of $\delta_{\text{PGI}:k}$, $r_k$, and $h_f^2$ given by $\hat{\delta}_{\text{PGI}:k}$, $\hat{r}_k$, and $\hat{h}_f^2$. A simple estimator of $k$ is

$$\hat{k}_0 = (1-\hat{r}_k)\,\frac{\hat{\delta}^2_{\text{PGI}:k} - \text{Var}\left(\hat{\delta}_{\text{PGI}:k}\right)}{\hat{h}_f^2},$$

where we subtract $\text{Var}\left(\hat{\delta}_{\text{PGI}:k}\right)$ to remove bias from sampling variation in $\hat{\delta}_{\text{PGI}:k}$. However, this estimator can still be substantially biased by noise in the estimate of $h_f^2$. The expectation of $\hat{k}_0$ is

$$\mathbb{E}[\hat{k}_0] = (1-r_k)\delta^2_{\text{PGI}:k}\mathbb{E}[(\hat{h}_f^2)^{-1}], \tag{59}$$

which we approximate with a second-order Taylor expansion:

$$\mathbb{E}[\hat{k}_0] \approx \left(1 + \frac{\text{Var}(\hat{h}_f^2)}{(h_f^2)^2}\right)k. \tag{60}$$

This yields a bias-corrected estimator of $k$:

$$\hat{k}_1 = \left(1 - \frac{\text{Var}(\hat{h}_f^2)}{(h_f^2)^2 + \text{Var}(\hat{h}_f^2)}\right)\hat{k}_0. \tag{61}$$

However, in real-world applications, we do not know the true value of $h_f^2$, so the denominator in the bias correction factor is unknown. Since $\mathbb{E}[(\hat{h}_f^2)^2] = (h_f^2)^2 + \text{Var}(\hat{h}_f^2)$, we propose the following bias-corrected estimator:

$$\hat{k} = \left(1 - \frac{\text{Var}(\hat{h}_f^2)}{(\hat{h}_f^2)^2}\right)\hat{k}_0 = \frac{(1-\hat{r}_k)(1-\hat{z}_f^{-2})(\hat{\delta}^2_{\text{PGI}:k} - \text{Var}(\hat{\delta}_{\text{PGI}:k}))}{\hat{h}_f^2}, \tag{62}$$

where

$$\hat{z}_f = \frac{\hat{h}_f^2}{\sqrt{\text{Var}(\hat{h}_f^2)}}. \tag{63}$$

### 9 Sampling variances of parameter estimates

Here we consider how to derive the sampling variances of the parameters estimated from the incomplete PGI (Figure 3). The estimating equations are non-linear functions of ($\delta_{\text{PGI}:k}$, $\alpha_{\text{PGI}:k}$), the direct effect and average NTC of the normalized incomplete DGE PGI on the normalized phenotype; $r_k$ the equilibrium correlation between maternal and paternal incomplete DGE PGIs; and $h^2_f = v_g/v^y_{\text{eq}}$. We assume that the sampling variance-covariance matrix of ($\delta_{\text{PGI}:k}$, $\alpha_{\text{PGI}:k}$) is known, and that the estimates of ($\delta_{\text{PGI}:k}$, $\alpha_{\text{PGI}:k}$), $r_k$, and $h^2_f$ are uncorrelated. This is obviously true when they are from independent data. However, even when they are derived from the same data, their correlations are unlikely to be high. This is because the estimate of $\delta_{\text{PGI}:k}$ derives from within-family variation, which is uncorrelated with the paternal and maternal PGIs from which the correlation $r_k$ is estimated. The estimate of $h^2_f$ could be correlated with $\delta_{\text{PGI}:k}$ when derived from the same data, but its correlation is probably not very high in most real-world scenarios.

Given the simplifying assumption of independence between the sample estimates, we can then obtain an approximation to the variance of some non-linear function $g\left(h^2_f, r_k, \delta_{\text{PGI}:k}, \alpha_{\text{PGI}:k}\right)$ by the Delta method:

$$\text{Var}\left(g\left(\hat{h}^2_f, \hat{r}_k, \hat{\delta}_{\text{PGI}:k}, \hat{\alpha}_{\text{PGI}:k}\right)\right) \approx \left(\frac{\partial g}{\partial \hat{h}^2_f}\right)^2 \text{Var}\left(\hat{h}^2_f\right) + \left(\frac{\partial g}{\partial \hat{r}_k}\right)^2 \text{Var}\left(\hat{r}_k\right) + \tag{64}$$

$$\left(\frac{\partial g}{\partial \hat{\delta}_{\text{PGI}:k}}\right)^2 \text{Var}\left(\hat{\delta}_{\text{PGI}:k}\right) + \left(\frac{\partial g}{\partial \hat{\alpha}_{\text{PGI}:k}}\right)^2 \text{Var}\left(\hat{\alpha}_k\right) + 2\left(\frac{\partial g}{\partial \hat{\alpha}_{\text{PGI}:k}}\right)\left(\frac{\partial g}{\partial \hat{\delta}_{\text{PGI}:k}}\right) \text{Cov}(\hat{\alpha}_{\text{PGI}:k}, \hat{\delta}_{\text{PGI}:k}). \tag{65}$$

The method implemented in *snipar* approximates the gradients numerically for each estimating function $g$ in order to calculate the approximate sampling variance for each parameter we estimate.

### 10 Simulation study of bias and sampling variance approximation

In order to investigate bias in our parameter estimation procedure and the accuracy of estimated standard errors, we simulated parameter inference in a scenario that mimics using the results of two-generation PGI analysis on $N$ independent trios and an unbiased estimate of $h^2_f$.

We set parameters to a scenario where we have AM at equilibrium and no indirect genetic effects, and where AM is due to matching on the phenotype in question. We set the equilibrium heritability to be $h^2_{\text{eq}} = 0.8$ and the phenotypic correlation between parents to $r_y = 0.75$, implying that $r_\delta = 0.8 \times 0.75 = 0.6$ and $h_f^2 = (1 - r_\delta)h_{\text{eq}}^2 = 0.32$. Given $k$, this implies that

$$\rho_k = 1 - (1-k)r_\delta;\ r_k = \frac{k r_\delta}{\rho_k};\ \delta_{\text{PGI}:k} = \sqrt{\frac{k h_f^2}{1 - r_k}};\ \beta = \frac{\delta_{\text{PGI}:k}}{\rho_k};\ \alpha_{\text{PGI}:k} = \frac{\beta - \delta_{\text{PGI}:k}}{1 + r_k}. \tag{66}$$

If we have estimated $\delta_{\text{PGI}:k}$ and $\alpha_{\text{PGI}:k}$ from a two-generation PGI analysis (with phenotype and PGIs standardized to have variance 1) using $N$ independent trios, then (see Equation 36)

$$\begin{bmatrix} \hat{\delta}_{\text{PGI}:k} \\ \hat{\alpha}_{\text{PGI}:k} \end{bmatrix} \sim \mathcal{N}\left( \begin{bmatrix} \delta_{\text{PGI}:k} \\ \alpha_{\text{PGI}:k} \end{bmatrix}, \frac{1 - R_{\text{trio}}^2}{N(1 - r_k)^2} \begin{bmatrix} 2(1 + r_k) & -(1 + r_k) \\ -(1 + r_k) & 1 \end{bmatrix} \right); \tag{67}$$

where $R_{\text{trio}}^2$ is the variance explained by regression onto proband and parental PGI, which is equal to

$$R_{\text{trio}}^2 = \delta_{\text{PGI}:k}^2 + 2(1 + r_k)\alpha_{\text{PGI}:k}(\alpha_{\text{PGI}:k} + \delta_{\text{PGI}:k}). \tag{68}$$

We obtain our simulation estimates of $\delta_{\text{PGI}:k}$ and $\alpha_{\text{PGI}:k}$ by simulating from the bivariate normal distribution given in (67). We obtain our estimate of $\beta_{\text{PGI}:k}$ as $\hat{\beta}_{\text{PGI}:k} = \hat{\delta}_{\text{PGI}:k} + (1 + r_k)\hat{\alpha}_{\text{PGI}:k}$.

We consider that we have estimated $r_k$ by computing the sample correlation coefficient between the parents' PGIs across the $N$ independent trios, implying that the approximate sampling distribution of $\hat{r}_k$ is

$$\hat{r}_k \sim \mathcal{N}\left( r_k, \frac{(1 - r_k^2)^2}{N} \right). \tag{69}$$

We simulated $\hat{r}_k$ independently from $\hat{\delta}_{\text{PGI}:k}$ and $\hat{\alpha}_{\text{PGI}:k}$ as we found in our other simulations of trio genotype data that estimates of $\hat{r}_k$ were uncorrelated with $\hat{\delta}_{\text{PGI}:k}$ and $\hat{\alpha}_{\text{PGI}:k}$ when estimated from the correlation between parents' PGIs.

We simulated estimates of $h_f^2$ from a normal distribution:

$$\hat{h}_f^2 \sim \mathcal{N}(h_f^2, \text{Var}(\hat{h}_f^2)), \tag{70}$$

and we varied the sampling variance between simulations. We did not analyze simulation replicates where $\hat{h}_f^2 < 0$.

We considered $N = 10^4$ and $N = 5 \times 10^5$, $k = 0.05, 0.2, 0.5$, and $\sqrt{\text{Var}(\hat{h}_f^2))} = 0.01, 0.1$. We considered all combinations of these parameters, giving 12 simulations in total. For each set of simulation parameters, we simulated 1000 replicates. For each set of parameters, we assessed the bias by comparing the sample mean of the estimated parameters across the replicates, and we assessed the estimated standard errors by comparing the median standard error estimate across the replicates to the standard deviation of the parameter estimates across the replicates. We give the results in Supplementary Table 7.

### References


[1] Crow, J. F. and Kimura, M. *An introduction to population genetics theory.* New York, Evanston and London: Harper & Row, Publishers 1970. Publication Title: The Blackburn Press.

[2] Young, A. I., et al. Mendelian imputation of parental genotypes improves estimates of direct genetic effects. *Nature Genetics*, **54**(6):897–905 2022. ISSN 1546-1718. doi:10.1038/s41588-022-01085-0.

[3] Nagylaki, T. Assortative mating for a quantitative character. *Journal of Mathematical Biology*, **16**(1):57–74 1982. ISSN 14321416. doi:10.1007/BF00275161.

[4] Visscher, P. M., et al. Assumption-free estimation of heritability from genome-wide identity-by-descent sharing between full siblings. *PLoS Genetics*, **2**(3):e41 2006. ISSN 1553-7404. doi:10.1371/journal.pgen.0020041.

[5] Young, A. I., et al. Relatedness disequilibrium regression estimates heritability without environmental bias. *Nature Genetics*, **50**(9):1304–1310 2018. ISSN 1546-1718. doi:10.1038/s41588-018-0178-9.

[6] Wright, S. Systems of mating. III. Assortative mating based on somatic resemblance. *Genetics*, **6**(2):144 1921. Publisher: Genetics Society of America.
